# An anti-steatosis response regulated by oleic acid through lipid droplet-mediated ERAD enhancement

**DOI:** 10.1101/2022.06.15.496302

**Authors:** Jorge Iván Castillo-Quan, Michael J. Steinbaugh, L. Paulette Fernández-Cárdenas, Nancy K. Pohl, Ziyun Wu, Feimei Zhu, Natalie Moroz, Veronica Teixeira, Monet S. Bland, Nicolas J. Lehrbach, Lorenza E. Moronetti Mazzeo, Magdalena Teufl, T. Keith Blackwell

## Abstract

Although excessive lipid accumulation is a hallmark of obesity-related pathologies, some lipids are beneficial. Oleic acid (OA), the most abundant monounsaturated fatty acid (FA), promotes health and longevity. Here we show that OA benefits *C. elegans* by activating the endoplasmic reticulum (ER)-resident transcription factor SKN-1A (Nrf1/NFE2L1) in a lipid homeostasis response. SKN-1A/Nrf1 is cleared from the ER by the ER-associated degradation (ERAD) machinery and stabilized when proteasome activity is low, and canonically maintains proteasome homeostasis. Unexpectedly, OA increases nuclear SKN-1A levels independently of proteasome activity, through lipid droplet (LD)-mediated enhancement of ERAD. In turn, SKN-1A reduces steatosis by reshaping the lipid metabolism transcriptome, and mediates longevity from OA provided through endogenous accumulation, reduced H3K4 trimethylation, or dietary supplementation. Our findings reveal a surprising mechanism of FA signal transduction, and a lipid homeostasis pathway that provides strategies for opposing steatosis and aging, and may mediate benefits of the OA-rich Mediterranean diet.

## Introduction

Lipids are vital for cellular functions and survival, and thus tight regulation of their levels and metabolism is critical. For example, the activation of the SREBP transcription factors at the ER membrane ensures the maintenance of minimum levels of cholesterol, phospholipids, and fatty acids (FAs)(*1, 2*). However, an obesogenic environment leads to a failure of lipid homeostasis and possibly to disease(*1, 3*). In the liver, for example, excess lipid accumulation (steatosis) can ultimately manifest as non-alcoholic fatty liver disease (NAFLD)(*3*). Given the current epidemic of obesity-related disease(*3, 4*), it is imperative to identify mechanisms that are linked to steatosis development or can protect against metabolic disturbance.

Work over the last few decades has revealed that lipid species vary widely in how they affect the organism. While an overload of saturated fatty acids (SFAs) is detrimental(*3, 4*), some mono- and polyunsaturated FAs (MUFAs and PUFAs respectively) are beneficial(*4, 5*). Notably, accumulation of the MUFA oleic acid (OA), a key component of the Mediterranean diet, can protect against lipotoxicity and disease(*3*). Among FAs, OA is the most potent inducer of lipid droplets (LDs), specialized organelles that bud off the ER membrane and store lipids(*6, 7*). While progress has been made in understanding how specific PUFAs act as signaling lipids(*4, 5*), the beneficial effects of OA have remained poorly understood. Humans, like most of the animal kingdom, cannot convert OA into PUFAs(*5*), suggesting that the beneficial effects of OA may be mediated by OA itself and not a metabolic derivative. It is a particularly intriguing question how this might occur because OA is a common dietary component and the most abundant FA within cells(*8*).

Various interventions that extend healthy lifespan in model organisms are associated with fat accumulation, including three examples in *C. elegans* involving OA(*9, 10*). Ablation of germline stem cells (GSCs) leads to accumulation of excess fat from unused yolk lipids(*11, 12*), elevated OA levels(*13*), and lifespan extension that depends upon OA production(*12, 14*). Inactivation of the COMPASS chromatin modification complex, which mediates trimethylation of lysine 4 on histone H3 (H3K4me3), also extends lifespan in an OA-dependent manner(*15*). In this paradigm the reduction in H3K4me3 activates SBP-1, the SREBP1 ortholog, which increases OA biosynthesis(*15*). Finally, dietary supplementation of OA is sufficient to extend *C. elegans* lifepan(*15, 16*). Each of these models provides an entry point for understanding how OA benefits the organism.

The transcription factor SKN-1 is required in various scenarios of *C. elegans* lifespan extension, including GSC ablation(*12*). GSC ablation increases SKN-1 activity dependent upon OA and the accompanying steatosis, through mechanisms that have remained unknown(*12*). The *skn-1* gene encodes two isoforms that have distinct functions and mammalian counterparts (see **Fig. S1a** for details)(*17, 18*). Like its ortholog NF-E2-related factor 2 (Nrf2/NFE2L2), the SKN-1C isoform is cytoplasmic and mediates an inducible antioxidant and xenobiotic defense response(*17, 19*). SKN-1A is the ortholog of the less well-characterized protein Nrf1 (NFE2L1), which maintains proteasome homeostasis(*18, 20*). Nrf1 is initially localized to the ER lumen and membrane, then is extruded from the ER by the ER-associated degradation (ERAD) machinery (**Fig. 1a**)(*20, 21*). ERAD is a multistep process that clears incompletely folded proteins from the ER and delivers them to the proteasome for degradation(*22, 23*). In its canonical model of activation, the proteasome recovery response, proteasomal insufficiency stabilizes Nrf1, which then increases proteasome subunit gene expression(*20*). This Nrf1 function and the features of its regulation are conserved in SKN-1A(*18, 24*). Interestingly, liver-specific Nrf1 knockout mice spontaneously develop NAFLD(*25, 26*), suggesting a role in lipid metabolism that has scarcely been explored. It has not been determined which of these SKN-1 isoforms mediate effects of GSC ablation. In this study, starting with an investigation of how GSC ablation increases SKN-1 activity, we have discovered a lipid homeostasis pathway in which OA activates SKN-1A/Nrf1 independently of proteasome activity by enhancing ERAD efficiency. In response, SKN-1A remodels lipid metabolism, reduces fat accumulation, enhances proteostasis, and extends lifespan.

**Fig. 1.**
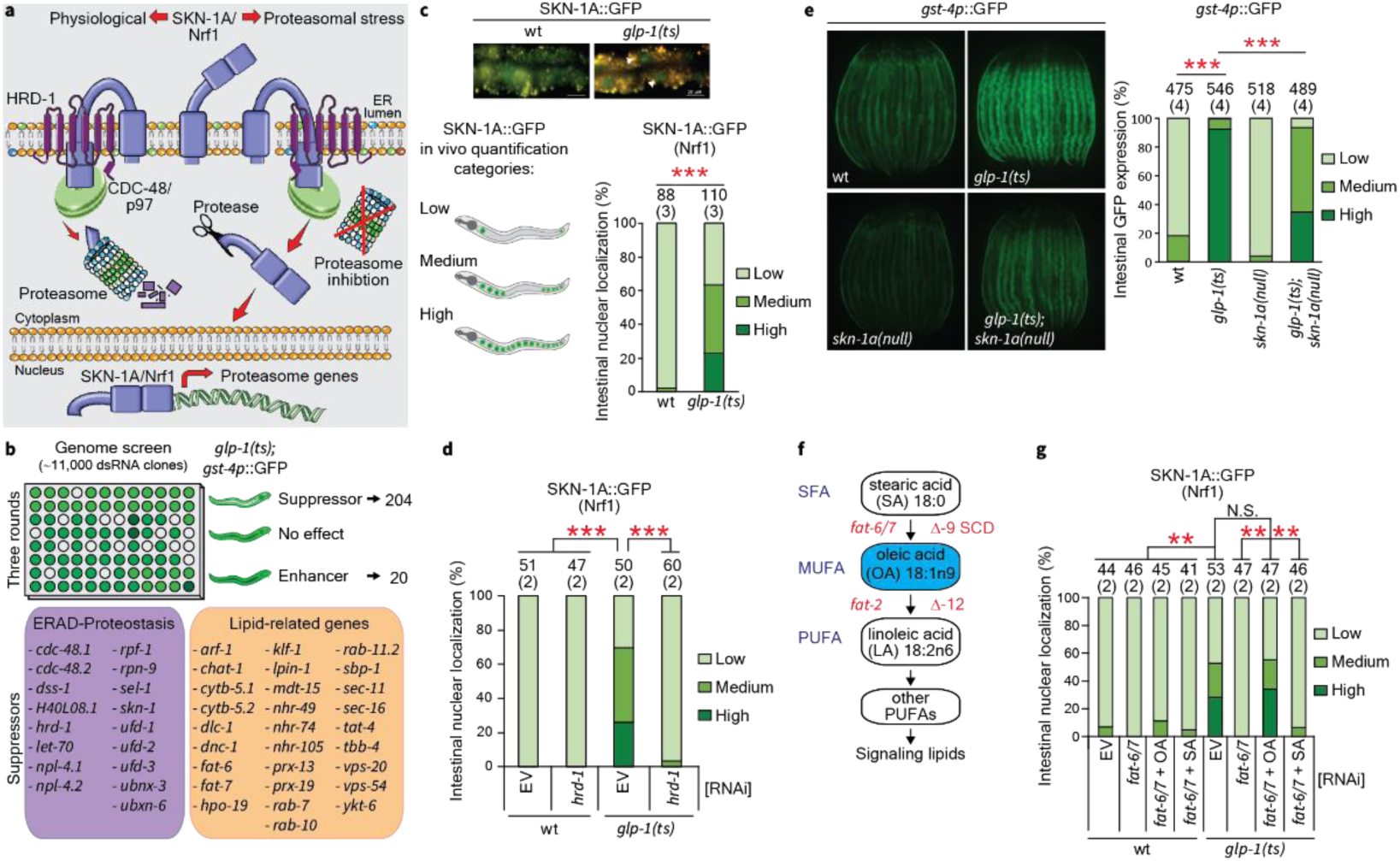
OA-dependent SKN-1A activation in GSC(-) *C. elegans*. **a**, Canonical proteasome recovery pathway mediated by SKN-1A/Nrf1(*18, 20*). HRD-1 and the CDC-48/p97 ATPase participate in the ERAD mechanism, which translocates SKN-1A/Nrf1 from the ER into the cytosol(*18, 20, 21*). **b**, A genome scale RNAi screen (see **Table S1** and **Supplementary Data 1** for details) identified ERAD and lipid-related genes as regulators of SKN-1 transcriptional activity in GSC(-) animals. We blocked GSC formation genetically, using a temperature-sensitive (ts) mutation in the *glp-1* (Notch) gene (*glp-1(ts)* in figures, see Methods)(*12*). Suppressor or enhancer refers to genes for which knockdown respectively decreased or increased reporter expression. **c**, SKN-1A nuclear accumulation (arrowheads) is increased in GSC(-) animals. Animals were classified (left) and quantified (right). **d**, SKN-1A nuclear accumulation in GSC(-) animals is dependent on the ERAD retrotranslocon HRD-1. **e**, Contribution of SKN-1A to *gst-4* expression in GSC(-) animals (see **Supplementary Data 2** for details). **f**, Abbreviated FA desaturation pathway (Δ-9 SCD: stearoyl-coA 9-desaturases; Δ-12: Delta-12-FA desaturase)(*30*). **g**, OA dependence of SKN-1A activation in GSC(-) animals. Unless otherwise indicated, all analyses were performed at day 1 of adulthood. Numbers above bars denote sample size (biological replicates). **p<0.01, ***p<0.001. Not significant (N.S. p>0.05).

## Results

### OA-dependent SKN-1A activation

To investigate how OA benefits the organism, we probed how GSC ablation increases SKN-1 activity by conducting a genome-scale RNA interference (RNAi) screen for effects on a SKN-1 target gene reporter (*gst-4p*::GFP) (**Fig. 1b**). We expected to identify SKN-1C/Nrf2 regulators because SKN-1C accumulates in nuclei in GSC-ablated (GSC(-)) animals(*12*), and because *gst-4* is known to be a SKN-1C target(*17*). Unexpectedly, however, ERAD-associated genes were prominent among our suppressor hits (**Fig. 1b, Table S1,** and **Supplementary Data 1**) suggesting involvement of SKN-1A/Nrf1 (**Fig. 1a**). Indeed, in GSC(-) animals transgenically encoded SKN-1A localized to nuclei in the intestine (**Fig. 1c** and **Supplementary Data2**), the liver and gut counterpart. SKN-1A nuclear localization depended on HRD-1 (**Fig. 1d** and **Supplementary Data 2**), the main component of the ERAD retrotranslocon channel(*22, 27*). By contrast, *hrd-1* knockdown did not reduce SKN-1C nuclear localization (**Fig. S1b** and **Supplementary Data 2**), linking the requirement for ERAD to SKN-1A. A mutation that specifically ablates *skn-1a*(*18*) (**Fig. S1a**) eliminated most expression of *gst-4p*::GFP (**Fig. 1e** and **Supplementary Data 2**), and RNAi against all SKN-1 isoforms abolished the remaining GFP signal (**Fig. S1c** and **Supplementary Data2**). Further indicating the importance of SKN-1A, it was required for the increased expression of another SKN-1C target (*gcs-1*(*28*)) in GSC(-) animals (**Fig. S1d**). The idea that SKN-1A and SKN-1C share regulation of these genes was unexpected but not surprising, because these SKN-1 isoforms share a common DNA binding domain (**Fig. S1a**). SKN-1A was not required for SKN-1C nuclear localization in GSC(-) animals (**Fig. S1e** and **Supplementary Data 2**), indicating that its importance does not derive from a secondary effect on SKN-1C. Together, the data suggest that the SKN-1 transcriptional response in GSC(-) animals is mediated predominantly by SKN-1A, with SKN-1C or other isoforms also contributing.

In GSC(-) animals SKN-1A nuclear accumulation was prevented by knockdown of the OA-synthesizing FA desaturases *fat-6/7* (**Fig. 1f-g** and **Supplementary Data 2**), indicating dependence upon OA. Further supporting this idea, dietary supplementation with OA but not the SFA stearic acid (**Fig. 1f**) relieved this block (**Fig. 1f-g** and **Supplementary Data 2**), and OA similarly rescued *gst-4p*::GFP activation (**Fig. S1f** and **Supplementary Data 2**). If OA activated SKN-1A through a PUFA intermediate, SKN-1A nuclear accumulation and activity should be blocked by knockdown of the FA desaturase FAT-2, which mediates the first step in converting OA into downstream PUFAs (**Fig. 1f**). By contrast, and arguing against a requirement for FAT-2, *fat-2* RNAi modestly increased SKN-1A activity (**Fig. S1g-h** and **Supplementary Data 2**). We conclude that in GSC(-) animals SKN-1A activation is stimulated specifically by OA, not downstream PUFAs.

### SKN-1A promotes proteostasis and longevity, and reduces steatosis

The extended lifespan of GSC(-) animals depends on the *skn-1* gene(*12*), but the isoform(s) responsible have not been identified. Consistent with its importance in the GSC(-) transcriptional response, SKN-1A was required for GSC(-) longevity (**Fig. 2a** and **Table S2**). Accordingly, knockdown of the ERAD retrotranslocon *hrd-1*, required for SKN-1A/Nrf1 activation(*18, 21*) (**Fig. 1d**), ablated the lifespan extension of GSC(-) animals and did not reduce their lifespan further in the *skn-1a* mutant background (**Fig. S2a and S2b** and **Table S2**). SKN-1A modestly extended lifespan when overexpressed (**Fig. S2c** and **Table S2**) and was fully or partially required in three other scenarios where the *skn-1* gene is needed for lifespan extension(*17*): dietary restriction (DR) and reduced activity of the growth and metabolism regulators mTORC1 (mechanistic target of rapamycin complex 1) or insulin/IGF-1 signaling (IIS) (**Fig. 2b-d** and **Table S2**). Thus, SKN-1A/Nrf1 is a broadly important modulator of aging, and a major contributor to lifespan extension effects that have previously attributed to the *skn-1* gene overall.

**Fig. 2.**
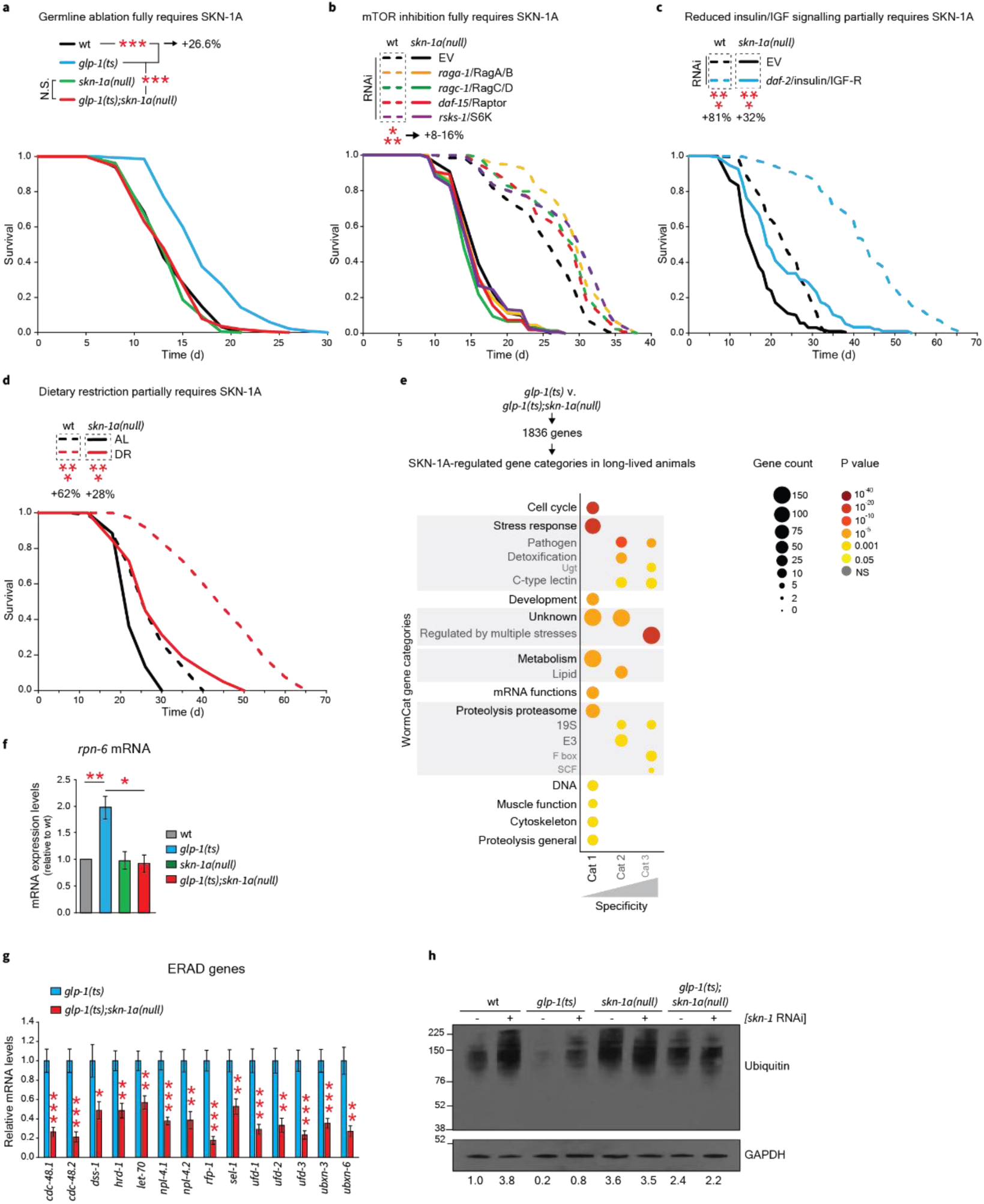
OA-dependent SKN-1A activity promotes longevity and enhances proteostasis. **a**, Lifespan extension from GSC ablation requires SKN-1A (see **Table S2** for replicates and statistics). **b**, Lifespan extension from knockdown of mTORC1 pathway components requires SKN-1A. **c-d**, Lifespan extension from *daf-2* knockdown (**c**) or DR (by food dilution) (**d**) partially requires SKN-1A. The lifespan of GSC(-) animals (**a**) was analyzed at 25°C, while others experiments (**b-d**) were run at 20°C. **e**, Gene categories modulated by SKN-1A specifically in GSC(-) animals. RNA sequencing (RNAseq) was analyzed with the gene annotation tool WormCat(*75*). We analyzed **Fig. 2e** and **Fig. S2b** separately because the germline accounts for about two-thirds of all adult nuclei(*12*), complicating direct comparisons between GSC(+) and GSC(-) transcriptomes(*12*). **f-g**, SKN-1A positively regulates the proteasome subunit gene *rpn-6* (**f**) and many ERAD genes (**g**), as measured by qRT-PCR and RNAseq respectively (shown as mean ± SEM; N=3; t-test). **h**, SKN-1A increases clearance of ubiquitinated proteins largely independently of other SKN-1 isoforms (quantification at the bottom reflect the average of two biological replicates of 3,000 animals each). Unless otherwise indicated survival experiments were initiated at the L4 stage (time 0). *p<0.05, **p<0.01, ***p<0.001.

To identify processes controlled by SKN-1A we used RNA sequencing (RNAseq) to investigate how ablation of *skn-1a* affected gene expression in wild type (WT) and GSC(-) animals (**Fig. 2e** and **Fig. S2d**). We focused our analysis on the much larger number of *skn-1a*-dependent genes that were identified in the long-lived GSC(-) background, in which SKN-1A is activated. For example, proteasomal subunit genes were upregulated in these animals (**Fig. 2e** and **Fig. S2e**), which exhibit increased proteasome gene expression and proteasome activity(*12, 29*). SKN-1A was required for expression of these proteasome subunit genes (**Fig. S2e**), including *rpn-6.1* (**Fig. 2f**), which has been linked to increased proteasome activity and longevity(*29*). Accordingly, SKN-1A was fully required for the remarkable resistance to the proteasome inhibitor bortezomib that is characteristic of GSC(-) animals (**Fig. S2f** and **Table S2**). Loss of *skn-1a* also reduced expression of all ERAD components that we identified in our screen (**Fig. 1b** and **Fig. 2g**), suggesting a multifaceted role in clearance of misfolded proteins. Consistent with these findings, GSC(-) worms exhibited a reduction in ubiquitinated proteins that was largely SKN-1A -dependent, with knockdown of other SKN-1 isoforms having no substantial additive effect (**Fig. 2h**). Thus, the SKN-1A isoform mediates the proteostasis benefits of GSC ablation.

Notably, lipid metabolism genes were enriched specifically among the SKN-1A-regulated genes detected in GSC(-) animals (**Fig. 2e**). Loss of *skn-1a* increased expression of lipid biosynthesis genes, most notably the transcription factor and master regulator *sbp-1*/SREBP1, as well as *dgat-2*/MOGAT1, and many FA desaturases and lipid transporters (**Fig. 3a**). By contrast, these *skn-1a* mutants exhibited decreased expression of LD biogenesis genes and the LD-associated TAG lipase *atgl-1* (**Fig. 3a**), which hydrolyses TAGs into FAs prior to β-oxidation(*30*) (**Fig. S3a**). SKN-1A also regulated expression of peroxisomal or mitochondrial β-oxidation enzyme genes (**Fig. S3b-c**). The *skn-1* gene has been implicated in lipid-related functions(*12, 31, 32*), but the isoform(s) involved have not been identified. In WT and GSC(-) animals loss of *skn-1a* led to increased intestinal fat accumulation (**Fig. 3b-c)** that was not elevated further by knockdown of other *skn-1* isoforms, suggesting that among SKN-1 isoforms SKN-1A may be unique in suppressing steatosis (**Fig. 3c** and **Fig. S3d**). Lack of *skn-1a* also led to increased LD size and number (**Fig. 3d-f)**. Thus, SKN-1A prevents excess lipid accumulation, possibly by reducing the expression of lipid biosynthetic enzymes and increasing aspects of lipid catabolism.

**Fig. 3.**
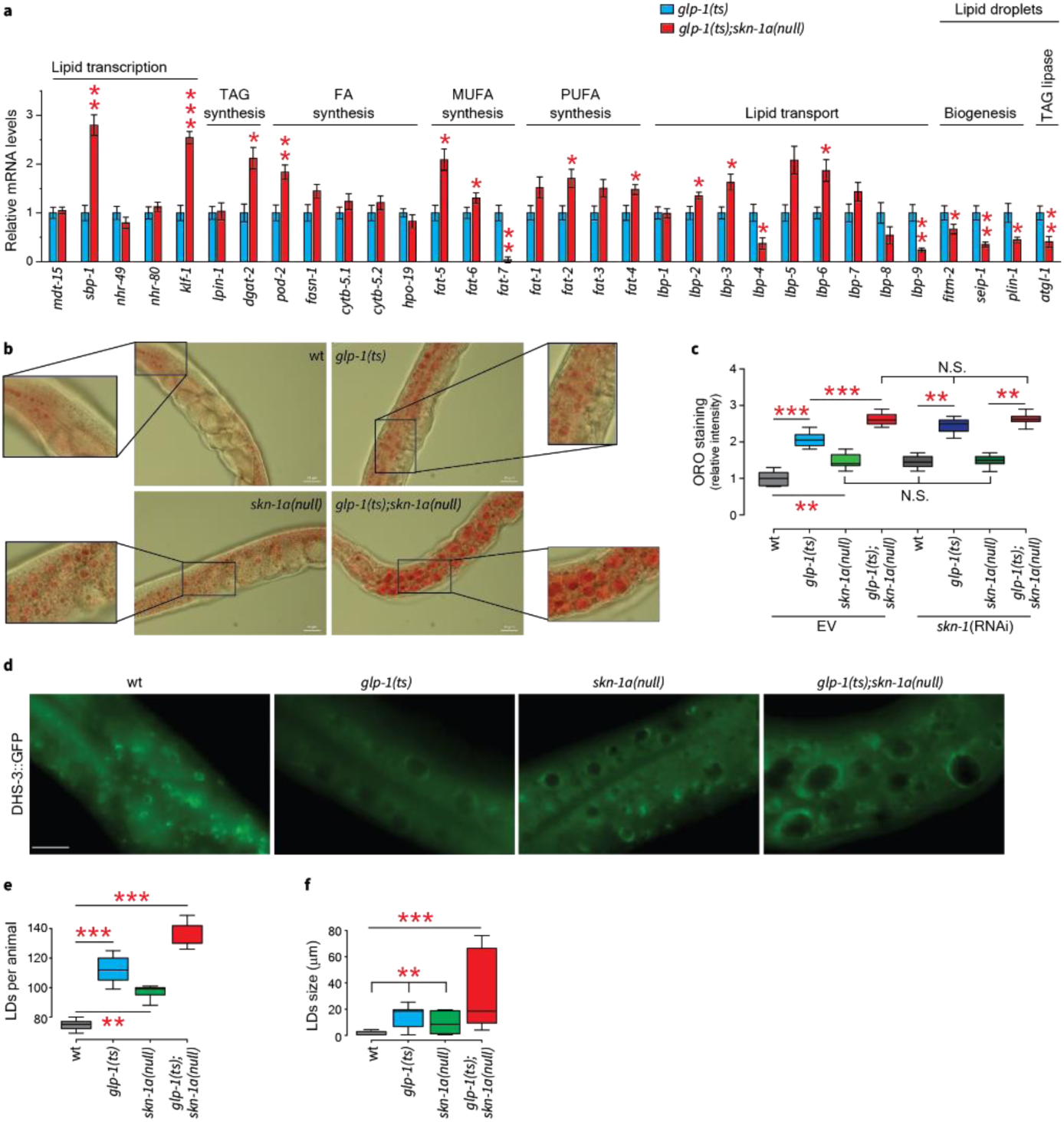
SKN-1A reduces fat accumulation in LDs. **a**, Regulation of lipid metabolism genes in GSC(-) animals by SKN-1A (RNAseq data shown; mean ± SEM; N=3; t-test). **b**, SKN-1A reduces TAG accumulation, visualized by fixed-ORO staining of 1-day old animals. **c**, SKN-1A mediates *skn-1* effects on TAG levels (quantification of **Fig. S3d**, one-way ANOVA, Tukey post hoc). **d-f**, SKN-1A reduces LD number (**e**) and size (**f**), visualized by the LD marker DHS-3::GFP(*77*). *p<0.05, **p<0.01, ***p<0.001. Not significant (N.S. p>0.05).

### OA-induced SKN-1A activation requires membrane homeostasis maintenance and LD functions

Our finding that in GSC(-) animals SKN-1A nuclear accumulation and activity are increased in an OA-dependent manner suggests that SKN-1A may respond to elevated levels of OA to reduce fat accumulation in a feedback loop. The only known paradigm for SKN-1A or Nrf1 activation is in the proteasome recovery response, in which these proteins are stabilized when proteasome activity is low (**Fig. 1a**)(*20, 24*). In GSC(-) worms the level of SKN-1A mRNA is not increased (**Fig. S4a**), suggesting that nuclear SKN-1A levels (**Fig. 1c**) are increased post-translationally. In this context SKN-1A nuclear accumulation and activity depend upon the ERAD machinery (**Fig. 1a** and **1d**), suggesting that SKN-1A is removed from the ER through the same set of mechanisms as in the proteasome recovery response. Remarkably, however, in GSC(-) animals proteasome gene expression and function and proteasomal stress resistance are dramatically elevated (**Fig. 2e-h**, and **Fig. S2e-f**)(*12, 29*). Thus, in GSC(-) animals nuclear SKN-1A levels are markedly increased by OA independently of proteasome activity, through a previously undescribed pathway that we refer to as the SKN-1A/Nrf1 lipid homeostasis response.

To investigate how OA-mediated SKN-1A activation occurs, we further examined results of genome scale and targeted RNAi screens involving GSC(-) animals. During processing SKN-1A is inserted into the ER membrane (**Fig. 1a**), the epicenter of lipid homeostasis(*2*). At the ER membrane OA, like other FAs, is either incorporated into membrane phospholipids or stored as TAGs within LDs(*6*) (**Fig. 4a**). We hypothesized that membrane phospholipid biosynthesis might be critical for OA-mediated SKN-1A activation. Phosphatidylcholine (PC), the most abundant ER and LD phospholipid(*2*), can be synthesized through the cytidine diphosphate diacylglycerol (CDP-DAG) pathway (**Fig. 4b**)(*30*). A screen of phospholipid synthesis enzymes (**Supplementary Data 1**) revealed that components of this pathway are required for SKN-1A to be activated by GSC ablation (*sams-1, pmt-1, pmt-2*; **Fig. 4c, Fig. S4b** and **Supplementary Data 2**) but not proteasome inhibition (**Fig. S4c-d** and **Supplementary Data 2**), suggesting that the CDP-DAG pathway is required specifically for OA-mediated SKN-1A activation. *C. elegans* depends upon PC synthesis through the CDP-DAG pathway because their laboratory diet lacks choline(*30, 33*), but with choline supplementation can synthesize PC when this pathway is inhibited(*30, 33*) (**Fig. 4d**). In GSC(-) animals that were supplemented with choline SKN-1A accumulated in nuclei independently of the CDP-DAG pathway (**Fig. 4c** and **Supplementary Data 2**), indicating a specific requirement for PC biosynthesis.

**Fig. 4.**
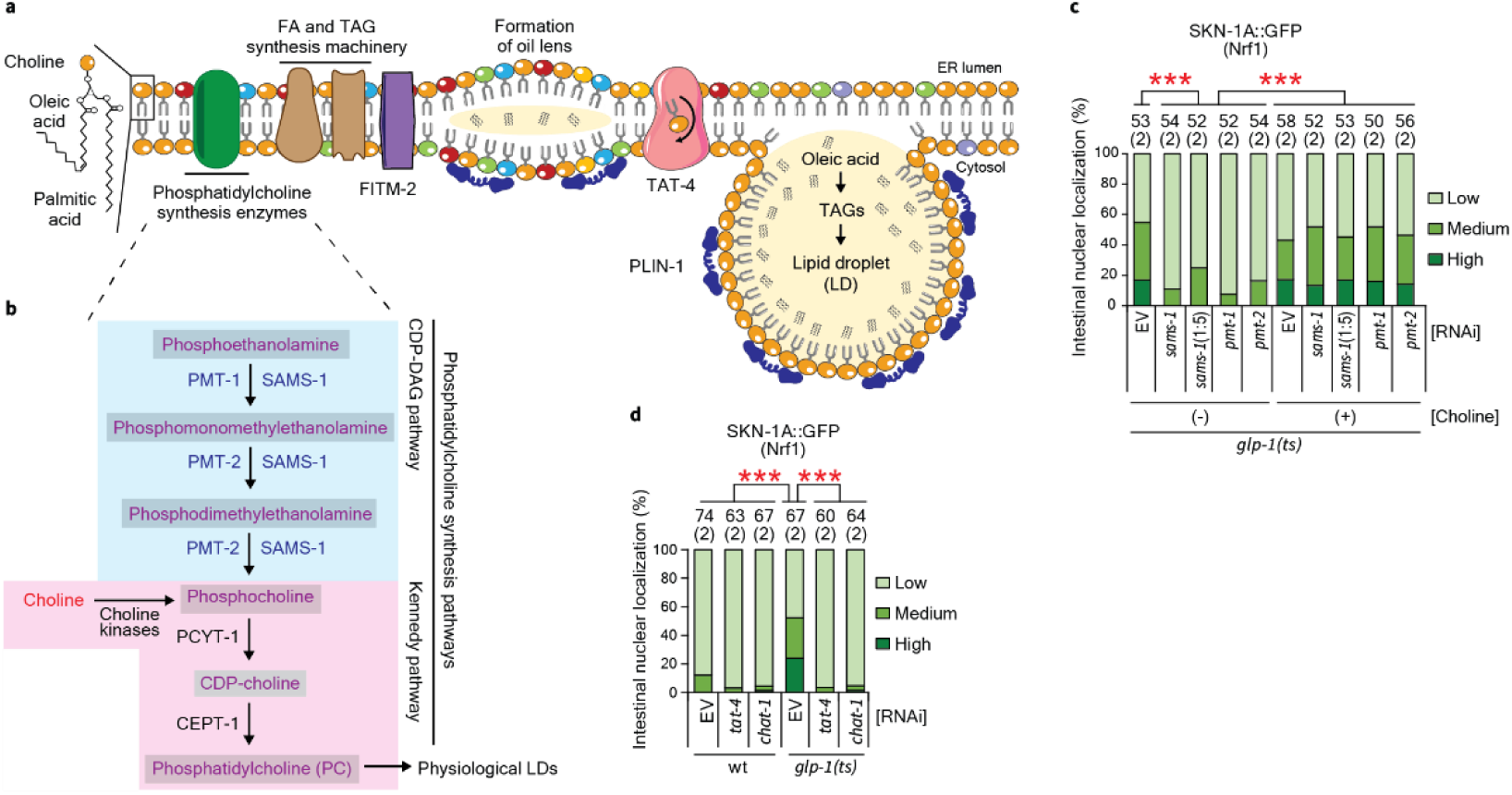
SKN-1A activation depends upon phospholipid membrane homeostasis. **a**, Diagram of ER membrane lipid homeostasis and LD formation mechanisms, highlighting those focused on here: 1) phosphatidylcholine (PC) biosynthesis enzymes (see **Fig. 4b**); 2) phospholipid flipping by TAT-4, ortholog of the transbilayer amphipath transporter (TAT), a P4 ATPase transmembrane flippase(*34, 37*); and 3) the ER transmembrane LD biogenesis factor FITM-2(*7, 41*) along with the LD membrane bound protein PLIN-1(*30, 77*). (see text and **Table S3** for more details). **b**, In *C. elegans* PC biosynthesis requires the PMT-1/2 methyltransferases and methyl groups generated by the *S*-adenosyl methionine synthase (SAMS-1)(*30*). **c**, SKN-1A activation in GSC(-) animals requires PC synthesis. Choline supplementation restores SKN-1A nuclear localization. *sams-1*(1:5) refers to a 1 in 5 dilution with EV used to avoid developmental delay (see Methods for details). **d**, SKN-1A activation in GSC(-) animals depends upon TAT-4 and its co-chaperone CHAT-1 (chaperonin of TAT-1; orthologous to transmembrane protein 30 (TMEM30))(*34, 37*) (see **Supplementary Data 2** for details). Numbers above bars denote sample size (biological replicates) ***p<0.001.

Newly synthesized phospholipids are incorporated into the cytosolic leaflet of the ER membrane and spontaneously equilibrate with the luminal side with the assistance of scramblases(*2, 34*). However the functions of the ER membrane in protein secretion and LD formation depend upon active membrane remodeling by phospholipid flippases(*34, 35*), which facilitate LD formation by increasing PC levels within the cytoplasmic leaflet (**Fig. 4a**)(*35, 36*). Our screen positives that suppressed *gst-4* expression (**Fig. 1b**) included the transmembrane ATP-dependent phospholipid flippase TAT-4 and its cofactor CHAT-1 (**Fig. S4e** and **Supplementary Data 2**), which move phospholipids from one side of the bilayer to the other (**Fig. 4a**)(*34, 37*). Knockdown of these genes also blocked SKN-1A nuclear accumulation in GSC(-) animals (**Fig. 4c**). Thus, OA-mediated SKN-1A activation depends upon both PC biosynthesis and active maintenance of membrane homeostasis.

OA is the most abundant FA within TAGs that are stored in LDs and the most potent inducer of LD formation(*6, 38*), and in GSC(-) animals the levels of OA(*13*) and fat storage in LDs (**Fig. 3b-f**) are elevated. Furthermore, OA-mediated SKN-1A activation depends upon membrane homeostasis and phospholipid production (**Fig. 4**), two parameters that influence LD biogenesis(*33, 39*). It was therefore intriguing that the suppressor hits from our screen (**Fig. 1b**) included enzymes essential for FA and TAG synthesis, the limiting and initial step of LD formation(*6, 40*), along with several proteins involved in LD budding or function (**Table S3**). Screening of other LD-related genes revealed additional suppressor hits (**Fig. 5a** and **Supplementary Data 1**) that were previously identified in *C. elegans* and human hepatic LD proteomes (**Fig. 5b**, **Fig. S5a**, and **Table S3**). Collectively, these LD-related screen hits included genes involved in TAG synthesis (*lpin-1*/LPIN) and LD expansion (*rab-7/10/11.2* and *arf-1)*, along with cytoskeleton components (*dlc-1, dnc-1, tbb-4*) associated with LD trafficking (**Table S3** for references, **Fig. 1b**, and **Supplementary Data 1**), as well as *fitm-2*, *plin-1*, and the peroxisomal β oxidation enzyme genes *dhs-28/*HSD17B4 and *daf-22*/SCP2 (**Fig. 5a**). FITM-2 is an ER transmembrane protein involved in segregating TAGs during formation of the oil lens for LD biogenesis (**Fig. 4a**)(*7, 41*). PLIN-1 is the worm ortholog of the perilipins, which have been linked to LD biogenesis and interactions with other organelles (**Fig. 4a**)(*6, 42*). Finally, LDs and peroxisomes interact physically and functionally (**Fig. S3a**), and share overlapping biogenesis mechanisms at the ER (**Table S3** for references). In *C. elegans* knockdown of *fitm-2* or deficiency of LD trafficking genes decreases LD biogenesis, knockdown of *plin-1* leads to clumped LDs, and inhibition of peroxisomal β oxidation or PC biosynthesis results in engorged LDs (**Table S3** for references). Our genetic manipulations led to similar effects, and knockdown of the phospholipid flippase *tat-4* reduced LD number and size (**Fig. 5c** and **Fig. S5c-d**). Thus, multiple mechanisms that are required for SKN-1A target gene activation have been implicated in LD biogenesis or functions.

**Fig. 5.**
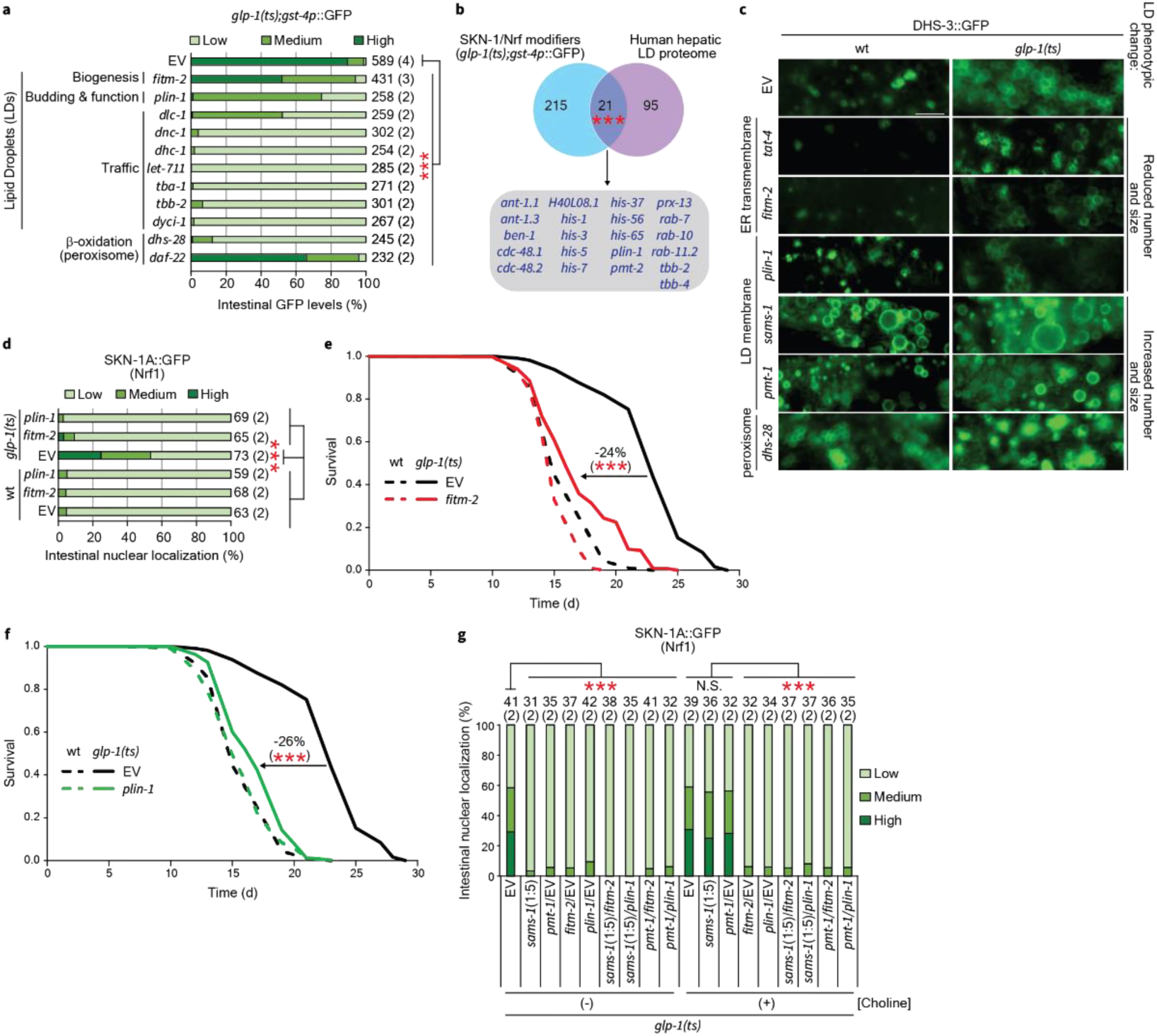
Dependence of OA-mediated SKN-1A activity on LDs. **a**, Suppressors of SKN-1A target (*gst-4*) activation in GSC(-) animals identified in a screen of LD-related genes (see **Table S2** and **Supplementary Data 1** for complete list and details). **b**, Overlap between *gst-4* suppressors and an LD proteome from human hepatocarcinoma cells(*79*) (see **Supplementary Data 3** for details). **c**, Genes required for SKN-1A activation alter LD morphology, as visualized by DHS-3::GFP. **d**, Knockdown of the LD genes *fitm-2* and *plin-1* blocks SKN-1A nuclear accumulation in GSC(-) animals (see **Supplementary Data 2** for details). **e-f**, Lifespan extension from GSC ablation requires the LD biogenesis/function proteins FITM-2 and PLIN-1 (see **Table S2** for details). **g**, SKN-1A activation requires *fitm-2* and *plin-1* independently of the CDP-DAG PC production pathway, revealed by choline rescue. Numbers above graph bars denote sample size (biological replicates), ***p<0.001. Not significant (N.S. p>0.05).

These observations led us to hypothesize that LD biogenesis or activities are required for OA to increase SKN-1A activity. Supporting this model, in GSC(-) animals knockdown of each LD-related screening suppressor hit that we tested reduced OA-dependent SKN-1A nuclear accumulation (*fitm-2*, *plin-1,* **Fig. 5d**, **Fig. S5d**, and **Supplementary Data 2**) and/or lifespan extension (*fitm-2*, *plin-1, dhs-28*, *daf-22,* **Fig. 5e-f, Fig. S5e-f,** and **Table S1**). The PC biosynthesis enzymes PMT-2 and SAMS-1 are present in LD proteomes (**Fig. 5b** and **Fig. S5a**), raising the possibility that LDs are needed for OA-induced SKN-1A activation because they might facilitate PC production (**Fig. 4b**). However, in GSC(-) animals choline supplementation failed to rescue SKN-1A nuclear accumulation when *plin-1* or *fitm-2* were knocked down simultaneously with these PC production genes (**Fig. 5g**, **Fig. S5g**, and **Supplementary Data 2**), indicating that OA-mediated SKN-1A activation and lifespan extension depend upon LD biogenesis and function *per se*.

### OA extends lifespan through the SKN-1A/Nrf lipid homeostasis response

Having established that in GSC(-) animals SKN-1A is activated through the SKN-1A/Nrf1 lipid homeostasis response (**Fig. S5h**), and that mechanisms mediating this response are essential for lifespan extension, we hypothesized that this would also be true in other scenarios where OA levels are elevated. In COMPASS-deficient animals, in which H3K4me3 formation is impaired, the resulting increase in OA production and levels drives lifespan extension(*15*). As our models predicted, in otherwise WT animals knockdown of the COMPASS methyltransferases *ash-2* or *set-2* activated the SKN-1A transcriptional response in an LD-dependent manner (**Fig. 6a, Fig. S6a-b** and **Supplementary Data 2)**. Knockdown of *ash-2* extended their lifespan, as shown previously(*15, 43, 44*), but shortened the lifespan of *skn-1a* mutants (**Fig. 6b** and **Table S2**), indicating that SKN-1A is required for lifespan to be extended by H3K4me3 impairment. Thus, the OA-induced lifespan extension that results from H3K4me3 impairment is mediated by the SKN-1A/Nrf1 lipid homeostasis response.

**Fig. 6.**
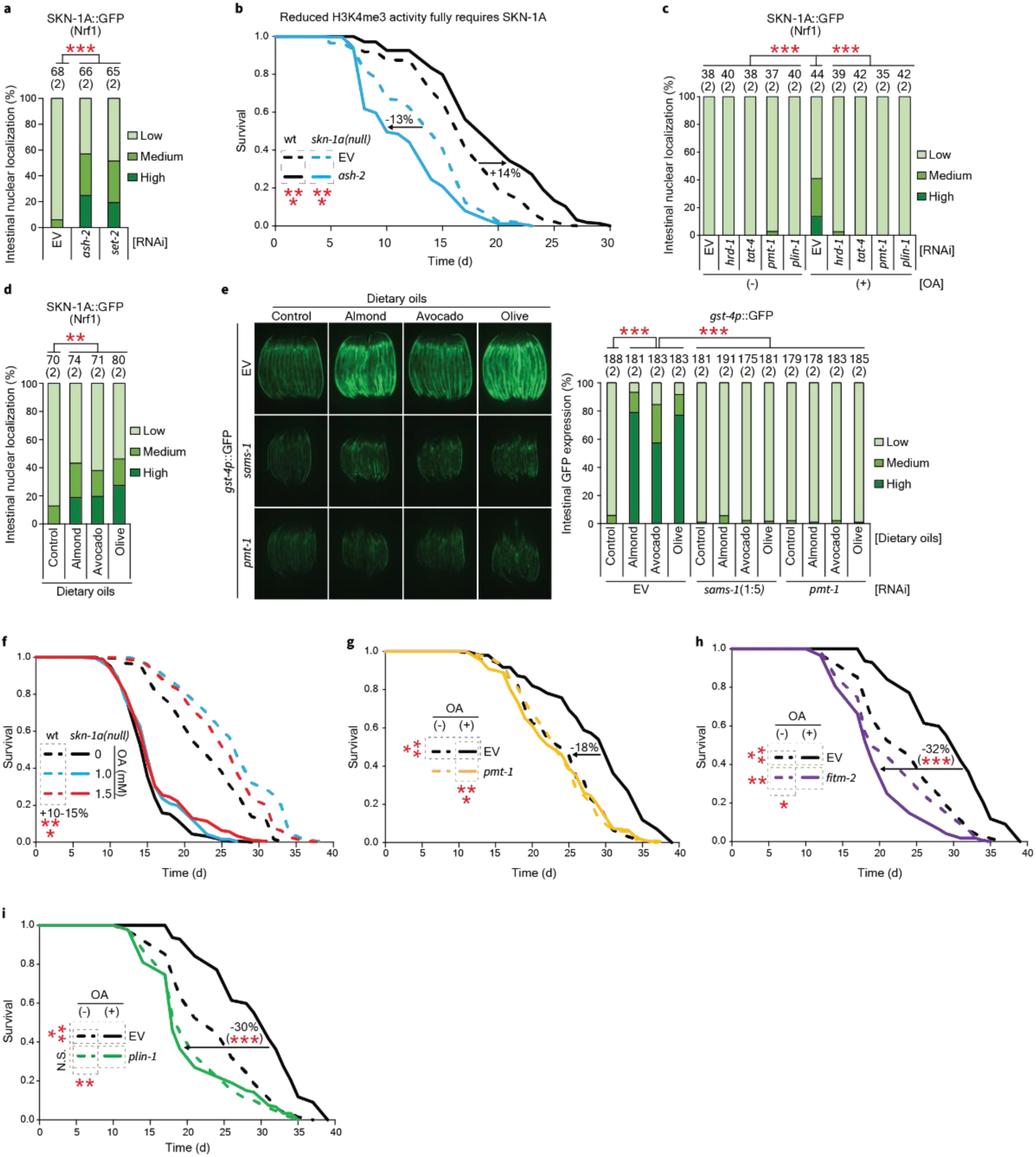
H3K4me3 deficiency and OA extend lifespan by enhancing SKN-1A activity through LDs and ERAD. **a**, Knockdown of H3K4me3 methyltransferases activates SKN-1A (see **Supplementary Data 2** for details). **b**, *ash-2* knockdown fails to extend lifespan in *skn-1a* null animals (see **Table S2** for details). **c**, SKN-1A is activated by OA feeding dependent upon ERAD (*hrd-1*), phospholipid flipping (*tat-4*), PC synthesis (*pmt-1*) and LD function (*plin-1*) (see **Supplementary Data 2** for details). **d**, OA-rich oils (see **Fig. S6e** for FA composition) activate SKN-1A in *C. elegans* (see **Supplementary Data 2** for details). **e**, Dietary oils increase *gst-4* expression in a PC-dependent manner (see **Supplementary Data 2** for details). **f**, Lifespan extension by OA requires SKN-1A (see **Table S2** for details). **g**, Lifespan extension by OA is blocked by knockdown of the PC-synthesizing enzyme *pmt-1* (see **Table S2** for details). **h-i**, The enhanced longevity of OA-supplemented animals is blocked by impairing LD formation or function (see **Table S2** for details). Numbers above bars denote sample size (biological replicates) *p<0.05, **p<0.01, ***p<0.001. Not significant (N.S. p>0.05).

Next, we investigated whether dietary supplementation with OA would be sufficient to activate SKN-1A through this pathway (**Fig. S5h**) and whether the resulting lifespan extension(*15, 16*) requires SKN-1A. In WT animals dietary OA increased SKN-1A nuclear accumulation dependent upon protein retrotranslocation through ERAD (*hrd-1*), membrane phospholipid homeostasis (*tat-4*), PC biosynthesis (*pmt-1*), and LD function (*plin-1*), and boosted *skn-1a*-dependent expression of the *gst-4p::GFP* target gene reporter (**Fig. 6c, Fig. S6c-d** and **Supplementary Data 2**). OA feeding thus activates SKN-1A through the SKN-1A/Nrf lipid homeostasis response. OA is the main MUFA component of olives and almonds, which are key components of the Mediterranean diet, as well as avocados (**Fig. S6e**)(*45*). As with OA supplementation, these OA-rich dietary oils increased nuclear SKN-1A and *gst-4* expression levels in a PC- and *skn-1a*-dependent manner (**Fig. 6d-e, Fig. S6f** and **Supplementary Data 2**). Thus, OA from various sources increases SKN-1A activity. Importantly, lifespan extension from OA supplementation depended not only upon SKN-1A (**Fig. 6f** and **Table S2**), but also upon genes involved in membrane homeostasis and LD-associated functions that we determined are required for its activation by OA (*pmt-1*, *plin-1*, *fitm-2*) (**Fig. 6g-i** and **Table S2**). We conclude that OA extends *C. elegans* lifespan by activating the SKN- 1A/Nrf1 lipid homeostasis response.

### LD-dependent ERAD enhancement by OA

Our results raise the intriguing mechanistic question of how OA activates SKN-1A even when proteasome activity is high (**Fig. 2e-h**). Given that OA induces formation of LDs(*6, 38*), which are essential for SKN-1A activation, the simplest model is that this activation is stimulated by LD formation, although it is also possible that membrane homeostasis mechanisms play a direct role. How might this increase SKN-1A levels? One possibility is OA increases SKN-1A retrotranslocation from the ER lumen to the cytosol by the ERAD machinery (**Fig. 1a**) without accelerating its degradation. ERAD involves a complex set of mechanisms in which different substrates may be processed by different overlapping components of the ERAD machinery(*22, 23*), and the precedent of many viral proteins that are extruded from the ER by ERAD components(*23*) illustrates that proteins can be retrotranslocated by the ERAD machinery without necessarily being degraded. Interestingly, links have been identified between ERAD and LDs: ERAD influences the LD proteome by degrading a subset of LD proteins, and some proteins involved in ERAD are present in LDs(*46, 47*). It has been proposed that LD formation might enhance the efficiency of ERAD(*47, 48*), although this unproven idea has remained controversial, in part because LD formation is not a generally essential prerequisite for ERAD(*49*). If OA simulates ERAD through LDs or other mechanisms that are required in the SKN-1A lipid homeostasis pathway, it would provide an attractive model to explain our findings.

To test this idea, we first investigated whether genes that are required for OA-induced SKN-1A activation might influence ERAD efficiency, using an ER luminal ERAD substrate reporter, mutated-CPL-1 (*nhx-2p*::CPL-1^W32A;Y35A^::YFP)(*50*). This substrate is identified in the ER lumen as a misfolded protein and targeted for ERAD-mediated degradation, and accumulates when ERAD is impaired (**Fig. S6g**). Importantly, levels of mutated-CPL-1 were increased by knockdown of each screening suppressor hit that we tested: the PC-biosynthesis enzymes PMT-1 and SAMS-1, the phospholipid flippase TAT-4, or the LD-associated proteins FITM-2 or PLIN-1 (**Fig. 7a-b**). Thus, ERAD was impaired in each case, indicating that baseline ERAD activity depends upon mechanisms that maintain ER membrane homeostasis or LD formation, and are essential for OA-mediated SKN-1A activation.

**Fig. 7.**
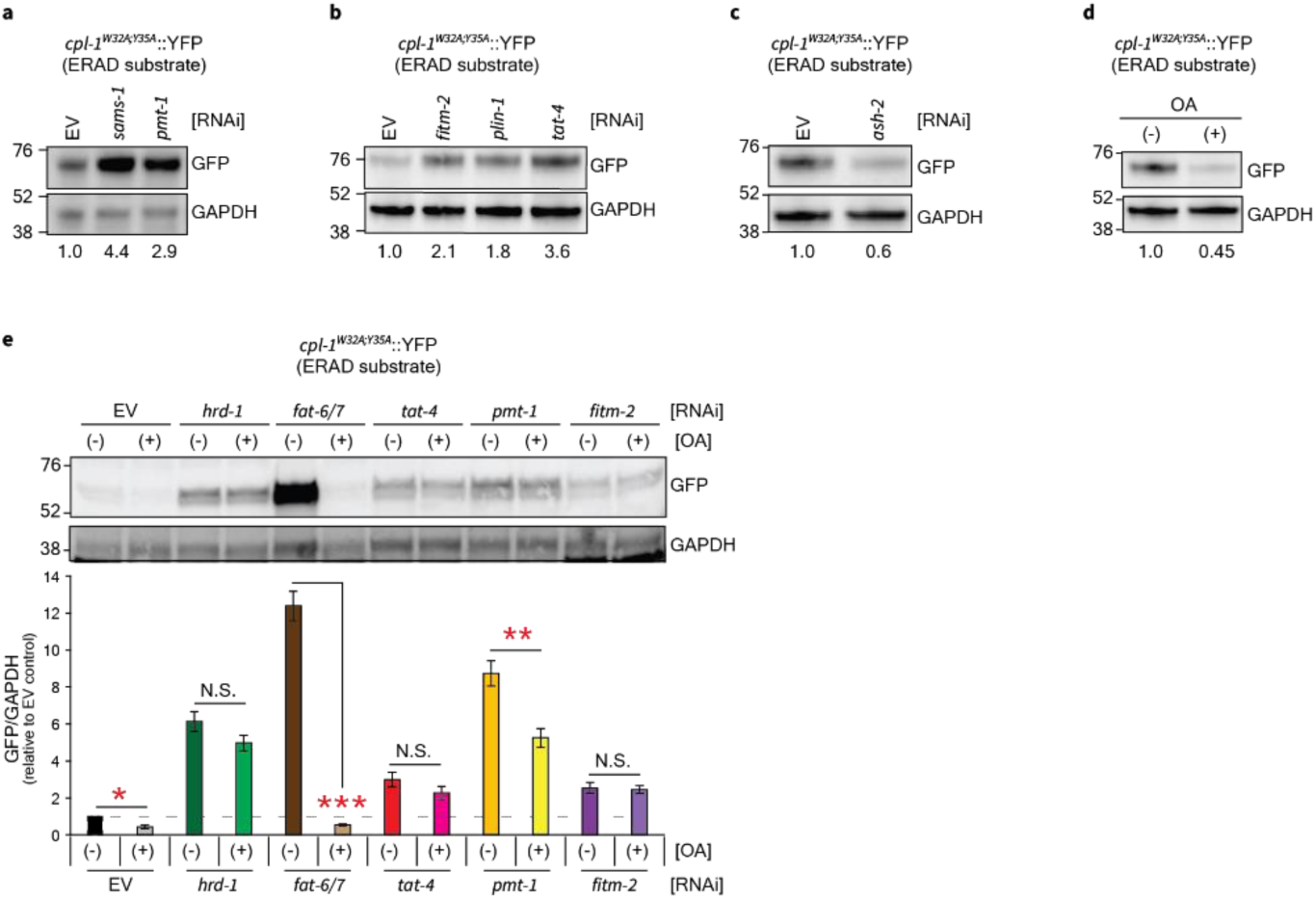
OA and LD homeostasis regulate ERAD efficiency. **a-b**, Impairment of ERAD by interference with PC biosynthesis (**a**), *tat-*4, *fitm-2,* or *plin-1* (**b**); quantification reflects the average of two biological replicates (3,000 animals each). **c**, Knockdown of the H3K4me3 COMPASS component *ash-2* enhances ERAD function (quantification at the bottom reflects the average of two biological replicates of 500 animals each). **d**, OA supplementation enhances ERAD function (quantification at the bottom reflects the average of two biological replicates of 3,000 animals each). **e**, OA-induced enhancement of ERAD (indicated by reduced levels of the mutated-CPL-1 substrate) is prevented by interference with the ERAD translocation channel (*hrd*-1), phospholipid flipping (*tat-4)*, or LDs (*fitm-2*). ERAD is impaired by interference with OA (*fat-6/7*) or PC synthesis (*pmt-1*), with robust rescue by OA feeding seen only with *fat-6/7* knockdown (data are presented as mean ± SEM; N=4 of 3,000 animals each; one-way ANOVA).

We next examined whether ERAD efficiency might be enhanced when OA levels are increased. If this is the case, then impairment of the COMPASS complex should facilitate ERAD. Accordingly, *ash-2* RNAi reduced the steady state levels of the mutated-CPL-1 ERAD substrate (**Fig. 7c**). This indicates that when H3K4me3 is reduced, ERAD is more efficient.

Remarkably, dietary OA supplementation was also sufficient to decrease the steady state levels of the ERAD substrate reporter (**Fig. 7d**). By contrast, ERAD was impaired by knockdown of the FA desaturases FAT-6/7 (**Fig. 7e**), which mediate the final step in OA biosynthesis (**Fig. 1f**). As would be predicted, this ERAD impairment was rescued by OA feeding (**Fig. 7e**). Together, these findings indicate that OA promotes protein clearance by ERAD and is critical for maintaining the normal level of ERAD activity. Importantly, disruption of active membrane homeostasis (TAT-4) or LD biogenesis (FITM-2) prevented OA supplementation from enhancing ERAD (**Fig. 7e**). Our data show that OA increases ERAD efficiency dependent upon LD- and membrane homeostasis-associated mechanisms that are required for it to increase nuclear accumulation of SKN-1A. Given that extrusion from the ER by ERAD is rate-limiting for SKN- 1A/Nrf1 processing and activity(*18, 20, 21*), the simplest model to explain our findings is that in the SKN-1/Nrf lipid homeostasis response OA increases SKN-1A activity by boosting ERAD efficiency, through its effects on ER membrane conditions and LD functions (**Fig. 8**).

**Fig. 8.**
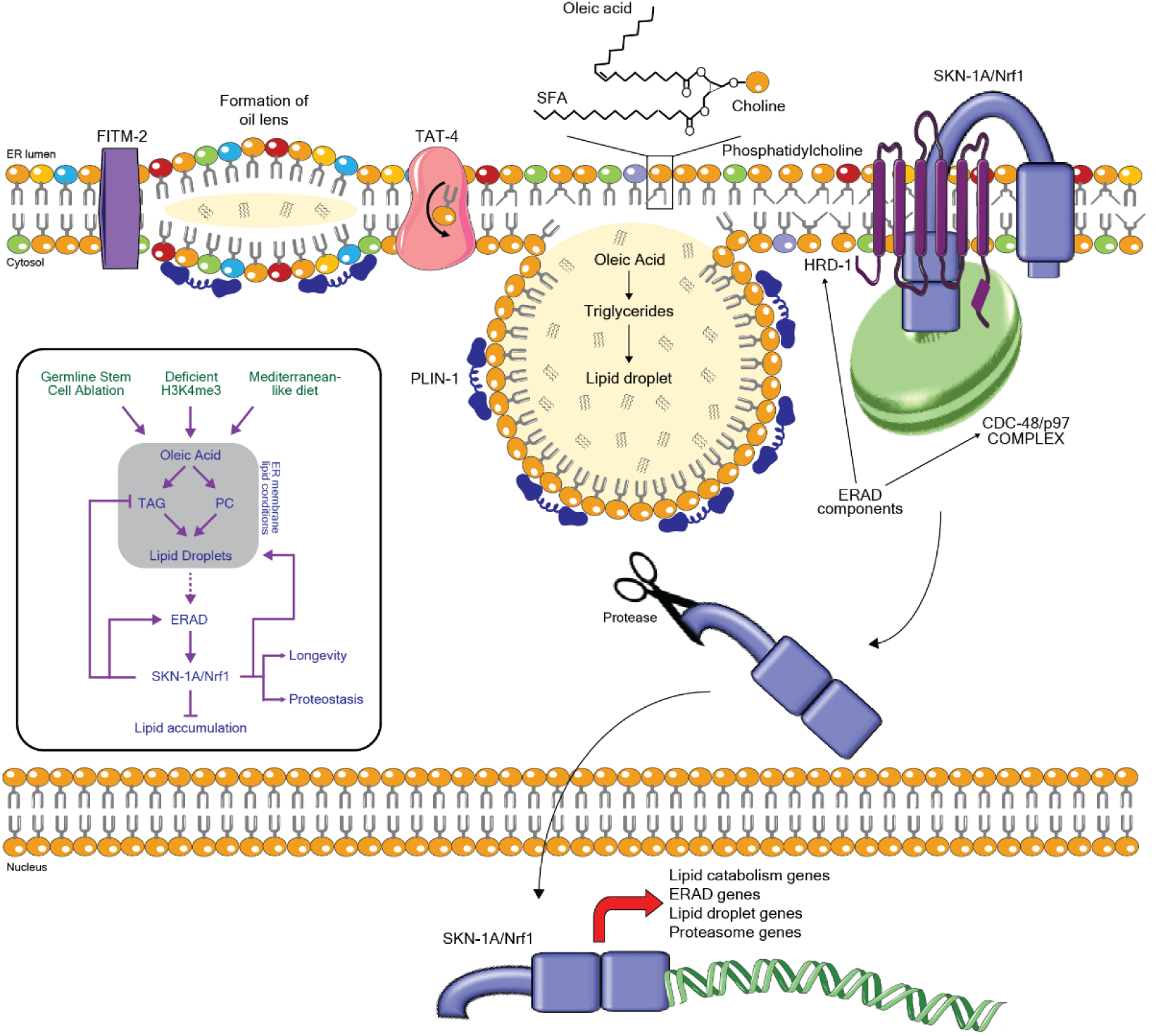
The SKN-1A/Nrf1 lipid homeostasis response. Model and summary diagram (inset) of the findings shown here. OA activates SKN-1A through a previously unknown mechanism that we term the SKN-1A/Nrf1 lipid homeostasis response. In this pathway, OA is sensed through LD formation and effects on ER membrane homeostasis. These changes enhance ERAD efficiency, thereby increasing levels of SKN-1A, which in turn remodels lipid metabolism gene expression and reduces fat storage in LDs. OA acts through this pathway to extend *C. elegans* lifespan.

## Discussion

An understanding of how specific lipids act as regulators is of great importance, given the current epidemic of diseases linked to excess fat storage and the promise of beneficial lipids for enhancing health and longevity(*9, 10, 45, 51*). Here we have determined that OA activates a homeostatic feedback loop in which it is sensed at the ER membrane to enhance ERAD efficiency, and thereby to activate SKN-1A (**Fig. 8**). In turn, SKN-1A stimulates a transcriptional program that remodels lipid metabolism, reduces lipid accumulation in LDs, promotes proteostasis, and extends lifespan. This SKN-1A/Nrf1 lipid homeostasis response pathway inhibits steatosis and is an essential mediator of the pro-longevity effects of OA, and may explain many of the remarkable health benefits of OA.

One of our most surprising findings is that OA is a major regulator of ERAD (**Fig. 7**). OA therefore promotes proteostasis through two mechanisms: by enhancing ERAD-mediated clearance of incompletely folded proteins from the ER and increasing proteasome gene expression through SKN-1A/Nrf1. OA could potentially affect ERAD in a variety of ways. OA is the most potent inducer of LD biogenesis(*6, 38*), and LD and membrane homeostasis functions are required for its effects on ERAD (**Fig. 7d-e**), suggesting the speculative but attractive model that OA promotes protein retrotranslocation through ERAD by inducing LD formation. One possibility is that LD formation or functions influence ERAD by altering ER membrane conditions such as lipid composition, packing, fluidity, or bending(*52, 53*). Biophysical interactions with the membrane bilayer are critical for HRD-1-mediated protein retrotranslocation(*27*), suggesting that membrane properties may be generally important for ERAD. Alternatively, crosstalk involving ERAD components and ERAD-regulated proteins that are present in the LD proteome may be critical(*46, 47*), or specific lipid metabolic products of LDs and/or peroxisomes might be important(*54*). Why would a regulatory relationship between OA and ERAD have evolved? It may be important that when lipid levels are high, OA could act as a sentinel that induces clearance of incompletely folded proteins, so that ER resources can be more effectively dedicated to lipid storage in LDs.

Another complex mechanistic question is how the OA-induced enhancement of ERAD increases SKN-1A activity. In the canonical proteasome recovery response, SKN-1A and Nrf1 are stabilized when proteasomal activity is reduced(*18, 20*), but in the SKN-1A/Nrf1 lipid homeostasis response OA increases nuclear SKN-1A levels even when proteasome activity is elevated (**Fig. S3** and **4b**). Why would OA increase ERAD-based elimination of an experimental substrate and not SKN-1A? It may be relevant that not all ERAD substrates are necessarily processed in the same way. In general, luminal and membrane-bound ERAD substrates are processed by overlapping but distinct sets of machinery(*22, 23*), and the paradigm that many viral proteins are retrotranslocated from the ER by different subsets of the ERAD machinery without degradation(*23, 48*) may also be instructive. Interestingly, in yeast a FAT-6/7-like desaturase that synthesizes OA is regulated by ER-membrane transcription factors (Spt23 and Mga2). When unsaturated FA levels are either high or low, these factors are subject to ERAD-dependent degradation or retrotranslocation and processing, respectively(*55*). This FA sensing is mediated through conformational changes within their transmembrane domains that are induced by differences in membrane lipid composition and packing(*55*). Thus, the relationship between OA and ERAD is ancient and provides an example of how membrane conditions can influence ERAD-mediated processing of a single client protein. Finally, it is also possible that SKN-1A is simply relatively resistant to proteasomal targeting, so that its levels can be elevated by increasing its retrotranslocation from the ER at a higher rate than it can be degraded.

SKN-1A/Nrf1 reduces fat storage in intestinal LDs (**Fig. 3c** and **Fig. S3d**), a function that is remarkably reminiscent of Nrf1 being required to prevent NAFLD in mice(*25, 26*). PC deficiency leads to steatosis and enlarged LDs in *C. elegans* and mammals(*33, 39*), and the most widely employed murine model of NAFLD is the methionine and choline deficient (MCD) diet, which impairs PC biosynthesis(*56*). SKN-1A activation depends upon PC synthesis, predicting that the MCD diet should not only alter LD formation but also should severely compromise Nrf1 and its capacity for reducing steatosis. Thus, Nrf1 is likely to be an important factor and promising therapeutic target in NAFLD prevention, a growing public health concern(*3*). The mammalian Nrf1 sequence contains a cholesterol recognition domain (not present in SKN-1A) through which cholesterol directly binds to Nrf1 at the ER membrane to inhibit its retrotranslocation(*57*). Evolution thus may have added a second dimension to Nrf1 regulation at the ER membrane, so that in mammals it receives both positive (OA/LD) and negative (cholesterol) signals.

Our finding that SKN-1A not only mediates OA-induced longevity but is also critical for lifespan extension in multiple settings (**Fig. 2b-d**) identifies it as an important modulator of aging. Given that the LD- and membrane-associated mechanisms we identified as being required for SKN-1A activation are also required for OA-induced lifespan extension, it may be that ERAD and SKN-1A are the convergent endpoint through which these LD-dependent effects of OA drive longevity, although it is certainly possible that LDs transduce the OA signal to increase lifespan through more than one mechanism. Besides altering lipid metabolism, OA-induced SKN-1A activation enhances proteostasis (**Fig. 2e-h** and **Fig. S2c-d**), a function associated with longevity and health(*17, 29*). It has been puzzling that GSC(-) and some other long-lived invertebrate models exhibit elevated fat storage(*9, 10*), a condition not typically associated with health(*3, 4*). Our results resolve this paradox in the models studied here, by showing that their elevated OA levels lead to SKN-1A activation. The importance of SKN-1 and Nrf2 for lifespan extension in *C. elegans* and *Drosophila*, respectively, has fueled interest in the role of Nrf2 in mammalian longevity(*17, 58*). However, Nrf2 is dispensable for the benefits of DR(*59*), raising the question of its overall importance. Our results indicate that Nrf1 should be examined as an important contributor to mammalian longevity and health.

SKN-1A can be activated by OA from various sources, including oils consumed by humans. Given the long life that OA confers upon *C. elegans*, and the evolutionary conservation from *C. elegans* to humans of the mechanisms that mediate the SKN-1A/Nrf1 lipid homeostasis pathway, it is tempting to speculate that Nrf1 may be a major mediator of the health benefits of olive oil and the Mediterranean diet. The mechanisms involved in the SKN- 1A/Nrf1 lipid homeostasis pathway, including ER membrane homeostasis, LD formation, and ERAD, may together represent a fruitful focus for development of therapies that reduce fat accumulation, prevent fatty liver disease, or slow aging.

## Materials and Methods

### *C. elegans* culture and maintenance

The wild-type N2 *C. elegans* strain was maintained at 20°C on nematode growth medium (NGM) plates seeded with *Escherichia coli* (OP50) using standard techniques(*60*). The GSC(-) model (or *glp- 1(ts)*) used corresponds to *glp-1(bn18)*(*12*). GSC(-) strains were maintained at 15 degrees (permissive temperature) and the following temperature shift protocol was used. Day-1 adult worms grown at 15°C were bleach-synchronized in 15 ml conical tubes. After washing the worms from the plates, animals were additionally washed for at least 3 times with M9 buffer to remove residual bacteria (centrifugation for washes was at 700 rpm for 1 min). After last wash the *C. elegans* pellet was resuspended in 6 mL of M9 buffer to which 650 μL of bleach and 1.3 mL of 5M NaOH were added. The tube was vortexed for 6 min after which the sample was washed at least 3 times with M9 buffer. The remaining *C. elegans* embryos were resuspended in 10 mL M9 buffer and left to hatch and arrest at the L1 stage for 24-36 hrs at 15°C. L1 animals were then seeded into the appropriate bacteria and placed again at 15°C. Animals were then shifted to the non-permissive temperature of 25°C (that does not allow development of the GCS) at the L2 stage and remained at this temperature for the duration of the experiment. Given the effect of temperature on many of the phenotypes we evaluated, non-GSC(-) strains were subjected to the same protocol when compared to GSC(-).

In all experiments, *glp-1(ts)* mutants were matched with the wild-type N2 strain used for outcrossing. In experiments that did not include GSC(-) strains, *C. elegans* were not subjected to a temperature shift protocol and instead were maintained at 20°C. After bleach-synchronizing, worms were starve-arrested at the L1 stage overnight in M9 buffer and seeded 12-15 hours later onto the corresponding bacteria. The *C. elegans* strains used in this study are detailed here:

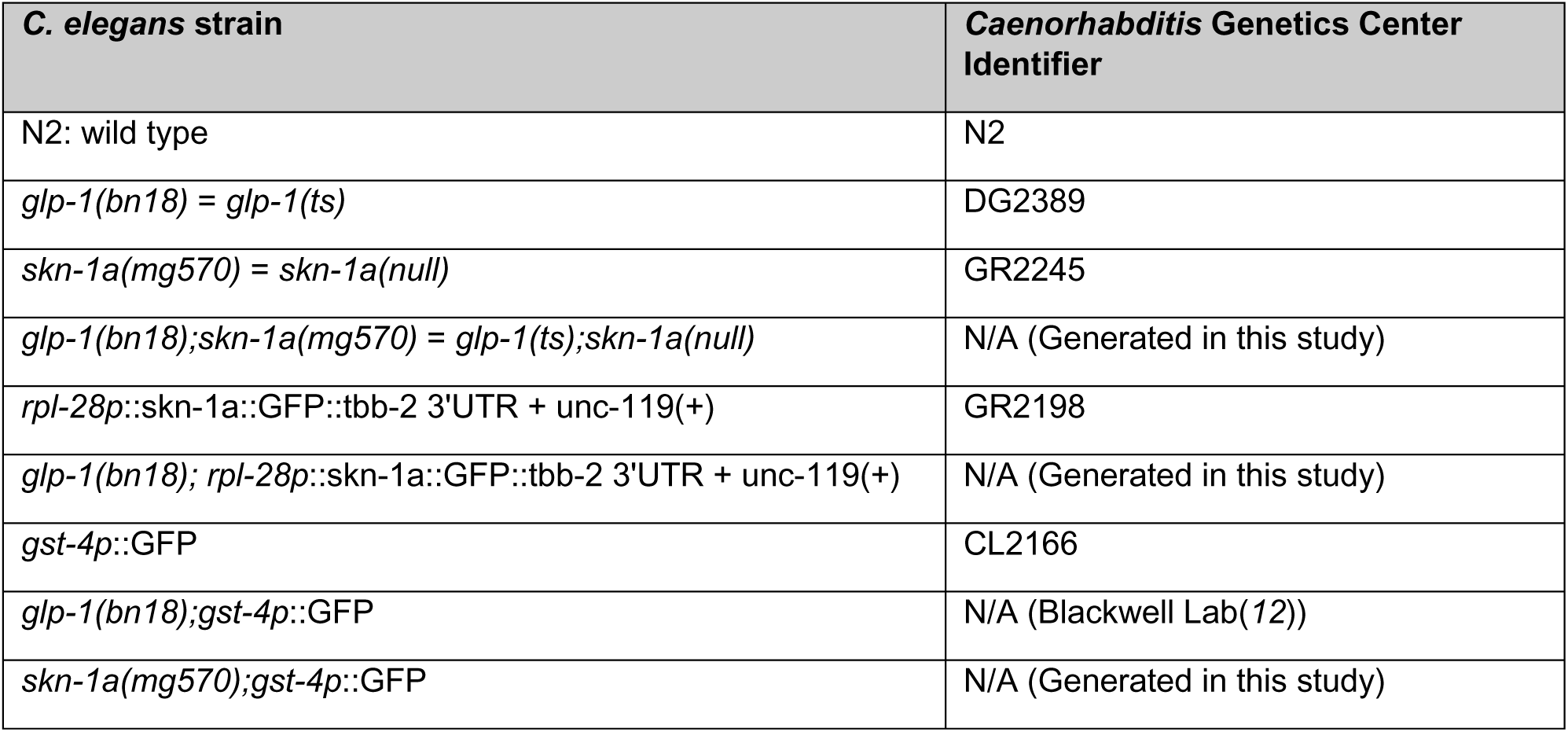
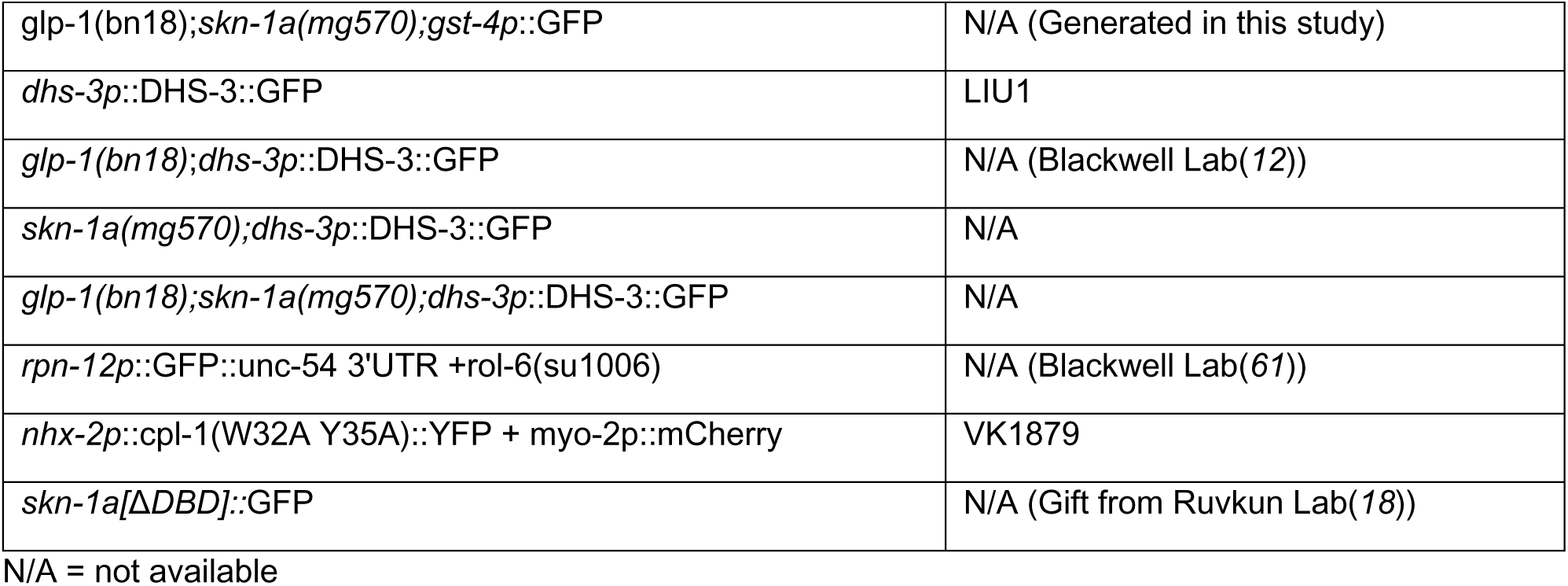

### *C. elegans* diet and RNAi

*C. elegans* were maintained on *E. coli* (OP50) bacteria as their standard food source. OP50 were cultured overnight in Luria-Bertani (LB) with 10 mg/L streptomycin at 37°C. 1 mL of the liquid culture was seeded on standard NGM plates (100mm x 15mm Petri dish) and air dried at room temperature before use. OP50 was also used in experiments in which RNAi was not performed. RNAi was performed by feeding *C. elegans* HT115 bacteria, using the pL4440 plasmid as empty vector (EV) control. The EV control was used in all experiments in which FA was supplemented to plates, even when RNAi was not performed. HT115 bacteria was grown using standard RNAi techniques(*62*). Briefly, RNAi cultures were grown overnight at 37°C with shaking at 220 rpm in 50 mL conical tubes with 5 mL LB medium containing 50 mg/mL carbenicillin and 50 mg/mL tetracycline. Cultures were then diluted 1:5 in LB containing the carbenicillin and tetracycline for 6 hours to allow for re-entry into the logarithmic growth phase. Cultures were then centrifuged at 4,500 rpm for 10 min and bacteria was concentrated to a volume of 5 mL LB containing 50 mg/mL carbenicillin, 50 mg/mL tetracycline and 0.2 g/L (1 mM) of IPTG prior to seed onto NGM plates containing 50 mg/mL carbenicillin, 50 mg/mL tetracycline and 0.2 g/L IPTG. For double RNAi, clones were grown separately in parallel and after spin-down equal amounts of two clones or one clone plus L4440 EV control were mixed and spread on plates. In cases were RNAi had to be diluted to allow *C. elegans* growth (example *sams-1*) the dilutant was the L4440 EV control. When a dilution was used this is shown in the figure, for example: *sams-1*(1:5) implies a 1 in 5 dilution. Bacterial clones were used in experimental analysis were sequence confirmed. RNAi was started at the L1 stage for all GFP reporter experiments, western blotting, RNAseq, RT-qPCT and TAG staining, except for experiments in which the H3K3me3 was inhibited which began at the embryonic stage (including lifespan). RNAi for lifespan experiments was initiated at the L4 stage, unless otherwise noted.

### Genome-scale and targeted RNAi screens

In our genome-scale RNAi screening of *glp-1(bn18);gst-4p*::GFP expression, we used the ORFeome (Vidal) library(*63*), and cherry picked libraries (gift from the Ruvkun laboratory) that include kinases, nuclear hormone receptors, and transcription factors derived from the Ahringer RNAi library(*62*). The genome scale screen was performed in duplicate and independently corroborated a third time. Clones that arrested or induced developmental delay were excluded during the genome-scale stage of the screen. Screen positives were grouped into functional categories (based on WormBase.org annotation) and the results were enriched by re-screening clones that classified in these functional groups from both the Vidal(*63*) and Ahringer RNAi libraries(*62*). At this stage clones that induced developmental abnormalities were diluted with the L4440 EV control to uncouple these phenotypes from direct effects on the SKN-1 transcriptional reporter. For example, the p97/*cdc-48* clones (*cdc-48* .1 and *cdc-48.2*) allowed normal development and growth rates at the 1:40 dilution. This allowed us to reintroduce these genes as suppressors that grouped in the ERAD-proteostasis category (**Fig. 1b, Table S1** and **Supplementary Data 1**). All clones that had an effect on GFP expression levels in the *glp-1(bn18);gst-4p*::GFP strain are shown in **Supplementary Data 1.**

The gene list for the phospholipid-targeted RNAi screen was obtained from a review describing lipid metabolism pathways(*30*). Genes were obtained from both the Ahringer and Vidal libraries(*62, 63*). The complete list of genes used for the phospholipid-targeted screen are described in **Supplementary Data 1**. The list of LD-related genes was initiated from those described in the aforementioned review and enriched by a literature search on PubMed.gov using the terms “elegans” and “droplet”. Genes reported to affect LD number and size, or that resided on the LD were added to the list. Genes that had already been analyzed for their effects on GFP expression levels on the *glp-1(bn18);gst-4p*::GFP strain in the previous screens were excluded for this list, for example: *pmt-1/2* that had been found to have an effect on *gst-4p*::GFP expression in the phospholipid-targeted screen. Genes were obtained from both the Ahringer and Vidal libraries(*62, 63*). The complete list of genes used for the LD-targeted screen (and references used to enrich the screen) are described in **Supplementary Data 1**.

### SKN-1/Nrf transcriptional GFP reporter analysis

The CL2166 strain *gst-4p*::GFP was used as a transcriptional reporter for SKN-1(*12, 64*). When used for analysis in experiments containing groups with *glp-1(ts),* animals in each group were maintained at 15°C and switched to 25°C at the L2 stage and remained at this temperature until scoring. L1 bleach-synchronized *C. elegans* were grown on the corresponding RNAi treatment. In experiments not containing *glp-1(ts)*, animals were maintained at 20°C throughout. For experiments in which the COMPASS H3K4me3 complex was knocked down animals were seeded at the embryonic stage and synchronized at the L4 stage. Animals were photographed and quantified at adulthood day 1. Scoring was as follows: low: no GFP or up to one third of intestinal cells; medium: up to two thirds of intestinal cells showed bright GFP or all intestinal cells showed dim GFP brightness; high: two thirds to all intestinal nuclei showed bright GFP. At least one trial of fluorescent reporters was scored blindly. All data presented in figures and supplemental figures are composites of at least two independent trials. In some experiments only the results with the GSC(-) line are shown in the figure to facilitate data visualization, but the wild type control groups were performed in parallel and can be found with all raw data in **Supplementary Data 2**. In figures, numbers above bars denote sample size (biological replicates). Chi-square test was used for statistical analyses using GraphPad Prism 9 (GraphPad Software, La Jolla, CA). For images, animals were immobilized for 5 min in 0.06% tetramisole/M9 buffer and aligned on non-seeded (empty) NGM plates. Images were taken with an Olympus IX51 microscope and cellSens standard 1.12 software.

### SKN-1A/Nrf1 nuclear localization

Animals were maintained at 15°C and synchronized at the L1 stage by bleaching. *C. elegans* were grown on the corresponding RNAi treatment or OP50 if no knockdown was performed, and kept at 15°C and switched to 25°C at the L2 stage where they were remained until day one adulthood for scoring. For *glp-1(ts)* the *rpl-28p*::skn-1a::GFP::tbb-2 was used, while for experiments not containing *glp-1(ts)* the *skn-1a*[Δ*DBD]::*GFP to facilitate visualization(*18*). For experiments not containing *glp-1(ts)*, animals were maintained at 20°C throughout. For experiments in which the COMPASS H3K4me3 complex was knocked down animals were seeded at the embryonic stage and synchronized at the L4 stage. On day one of adulthood animals were immobilized for 5 min in 0.06% tetramisole/M9 buffer, mounted on 2% agarose pads on glass slides under coverslips, and imaged with ZEN 2012 software on an Axio Imager.M2 microscope (Zeiss, Jena, Germany). Scoring was as follows: low: no GFP or up to one third observed in nuclei; medium: up to two thirds of nuclei showed GFP; high: two thirds to all intestinal nuclei showed GFP. In some experiments only the results with the GSC(-) line are shown in the figure to facilitate data visualization, but the wild type control groups were performed in parallel and can be found with all raw data in **Supplementary Data 2**. In figures, numbers above bars denote sample size (biological replicates). Chi-square test was used for statistical analyses using GraphPad Prism 9 (GraphPad Software, La Jolla, CA). At least one trial of these fluorescent reporters was scored blindly.

### Oil-red O TAG staining

Oil-red O (ORO; Sigma: O0625-25G) staining was used to stain neutral lipids, of which *C. elegans* possesses only TAGs(*11, 30, 65*). ORO staining was performed on fixed animals as previously described(*11, 12*), with some modifications. 500–1,000 L1 bleach-synchronized *C. elegans* were grown to adulthood day one and washed three times with phosphate-buffered saline (PBS). Upon the last wash, animals were treated with PBS containing 2% paraformaldehyde (PFA; Santa Cruz: SC-281692) then snap frozen in a dry ice/ethanol bath for at least 3 times to permeabilize the cuticle. *C. elegans* samples were then washed with PBS to remove the PFA at least twice with a 60% isopropanol solution. After removal of the isopropanol solution, samples were treated with freshly prepared and filtered ORO solution (0.5 g of ORO powder in 100 ml of 60% isopropanol). Samples were then stained for 1-3 hours in 1.5 ml Eppendorf tubes at room temperature with gentle shaking. Longer staining periods, such as overnight incubation(*11*), saturated ORO staining in *glp-1(ts)* animals to a level that rendered *glp-1(ts)* and *glp- 1(ts);skn-1a*(null) strains indistinguishable from each other. Animals were imaged at 40× using differential interference contrast microscopy. Quantification of ORO staining was performed on the upper to mid intestine. Statistical analysis (one-way ANOVA with Tukey post hoc analysis) was performed with GraphPad Prism 9.

### Lipid droplet quantification

Bleach-synchronized day-1 adult DHS-3::GFP *C. elegans* strains were grown as described above for GFP quantifications. In cases where RNAi was used, it was initiated after L1 arrest. All scoring was performed using day one adults. On day one of adulthood animals were immobilized for 5 min in 0.06% tetramisole/M9 buffer, mounted on 2% agarose pads on glass slides under coverslips, and imaged with ZEN 2012 software on an Axio Imager.M2 microscope (Zeiss, Jena, Germany). Quantifications were performed by scoring the droplets on images taken from 4 different regions of the *C. elegans* intestine of between 10-20 animals. The proximal and distal intestine and the mid region immediately before and after the gonad were used for analysis. The average of these regions was then plotted and analyzed using one-way ANOVA (with Tukey post hoc analysis) on GraphPad Prism 9.

### Longevity assays

Longevity experiments in *C. elegans* followed standard protocols(*12, 66*). Lifespan assays involving GSC(-) animals were performed at the non-permissive temperature of 25C, and all other lifespan experiments were performed at 20C. For all lifespan experiments *C. elegans* were L1 bleach-synchronized and seeded onto OP50 plates. Animals were transferred at mid to late L4 stage to fresh plates containing FUdR at a concentration of 50 μM to inhibit progeny development(*67*). For knockdown experiments, bacteria were prepared as described above and *C. elegans* were transferred at the L4 stage. For assays in which the COMPASS complex was inhibited *C. elegans* were seeded at the embryonic stage onto RNAi-containing plates and synchronized at the L4 stage by gonad developmental morphology. No FUdR was used for COMPASS experiments, in which animals were transferred every day. OA or choline supplementation for longevity experiments was initiated at the L4 stage. For *glp-1(ts)* experiments the growth protocol and temperature shift was as described above. *C. elegans* were maintained at a density of 25-35 *C. elegans* per 6 cm plate on live bacteria and scored every other day. Animals that crawled off the plate, ruptured, or died from internal hatching were censored. Lifespans were graphed as Kaplan–Meier survival curves and analysis of survival curves was generated using log-rank test as described(*68*). Additional statistical analysis was performed with GraphPad Prism 9 (GraphPad Software, La Jolla, CA). Key survival experiments were scored by multiple different people. All longevity assays shown in figures are composites of at least 2 independent biological replicates.

Details and statistics of all survival experiments are shown in **Table S2**.

### Proteasomal stress

Proteasomal stress was induced by treatment with the proteasome inhibitor bortezomib, (LC laboratories: B-1408) which was directly added io the NGM during plate pouring at 200 μM (stock diluted in DMSO at 50 mM). Plates were otherwise prepared and seeded with OP50 bacteria as described. *C. elegans* were transferred to these plates at the L4 stage and scored daily. Survival assays were graphed as Kaplan–Meier survival curves as previously described(*68*). Survival curve analysis were generated using log-rank test. All longevity assays shown in figures are composites of at least 2 independent biological replicates. Details and statistics of all survival experiments are shown in **Table S2**. For induction of proteasomal stress and analysis of proteasome subunit transcriptional reporters, animals were stimulated by adding 3 mL of bortezomib diluted in M9 buffer directly in the plates in which the animals were grown. GFP imaging was performed 24 hours later, and images were taken with Olympus IX51 microscope and cellSens standard 1.12 software.

### Micro- and macronutrient supplementation

Oleic and stearic acid (Sigma: 07501-1G and S3384-5G, respectively) supplementation was performed as previously described with some modifications(*69*). To ensure solubilization of the fatty acids (FAs), the NGM was supplemented with 0.001% Nonidet p40 substitute immediately after the media came out of the autoclave. This was done for both supplemented and control plates (a similar protocol was followed for the supplementation of oils). A FA stock solution (in double distilled and autoclaved water) was generated at 100 mM. After the solution of FAs (vehicle was used to adjust for volume in case different volumes of FA stock solution was used to adjust concentration and in the case of control plates) was added, the media was stirred for at least 40 min (under heating conditions that allowed the media to remain at 50°C) before pouring into the appropriate petri dishes. The most effective concentration of oleic acid was determined by three independent experiments in which 0.5, 1 and 1.5 mM were analyzed for activation of the SKN-1 transcriptional reporter *gst-4p*::GFP. In experiments in which only one concentration was used, the most effective one of 1 mM was used. The appropriate bacteria were seeded the day after and used within a week. For reporter analysis FA supplementation was initiated after L1 arrest, while for longevity assessment it began at the L4 stage.

For choline (Sigma C7527-100G) supplementation we used a stock solution of 3 M diluted in double distilled and autoclaved water. The final concentration used in the supplementation experiments was 30 mM added to the NGM as previously described (*33, 70*). Equal volume of vehicle was supplemented to control plates. During the optimization stages we compared the effect of 30 mM choline supplementation to 30 mM NaCl without observing any potential osmotic effects.

Dietary oils were supplemented to 1% in the NGM that contained 0.001% NP40 (Sigma: 74385- 1L) prepared as described above. Oils were obtained from Whole Foods: extra virgin olive oil and almond oil were the 365^TM^ Everyday Value brand, and avocado oil the Spectrum brand.

### Western blotting

Between 3-4,500 day-1 adults were used in each independent biological sample. Animals were synchronized by L1 arrest except in *ash-2* knockdown experiments, in which 500 animals were synchronized at the L4 stage by gonad developmental morphology. Animals were collected by washing plates with M9 buffer for at least 3 times. After removal of the M9 buffer in the last wash, the worm pellet was resuspended in lysis buffer (supplemented with protease and phosphatase inhibitors (Roche)) and sonicated at 30% for 13 seconds. The samples were then centrifuged at maximum velocity in Eppendorf tubes and supernatant was collected for BCA protein quantifications. Equal amounts of protein were eluted in 2 x sample buffer, boiled to 95°C, centrifuged at maximum velocity and loaded into NuPAGE Novex Bis-Tris 10% Gels. Antibodies used were ubiquitin (Abcam: ab7254; 1:500), GAPDH (Bethyl laboratories: A303-878A; 1:500), GFP (Sigma: SAB4301138; 1:500). All immunoblots are representative of at least two independent biological samples and average of their quantifications are shown under the blot images. Where 3 or 4 biological replicates were obtained the quantification and statistics are shown in bar-charts under the blot images.

### qRT-PCR

Samples were prepared from 1,500 day-1 adult bleach-synchronized *C. elegans* as previously described (*12*). Briefly, RNA was extracted using TRIzol-based (Fisher: 15596026) phenol-chloroform extraction and purified with RNA Clean and Concentrator-5 spin columns (Zymo research: R2050). RNA concentration and quality was assessed with a NanoDrop 1000 spectrophotometer (Thermo Fisher). cDNAs were prepared using SuperScript III First-Strand Synthesis SuperMix for qRT-PCR (Thermo Fisher). mRNA levels were quantified from biological triplicates and technical duplicates using SYBR Green (Thermo Fisher: 11760500) fluorescence on a 384-well format Real-Time PCR 7900 (Applied Biosystems, Foster City, CA). After an initial denaturation step (95°C for 10 min), amplification was performed using 40 cycles of denaturation (95°C for 15 s) and annealing (60°C for 1 min). Samples were analyzed by the standard curve method, with normalization to the reference genes *cdc-42* and *act-1*(*64, 71*). p values were calculated by t-student or one-way ANOVA (Tukey post hoc analysis) in GraphPad Prism 9. The primers used in this study are listed here:

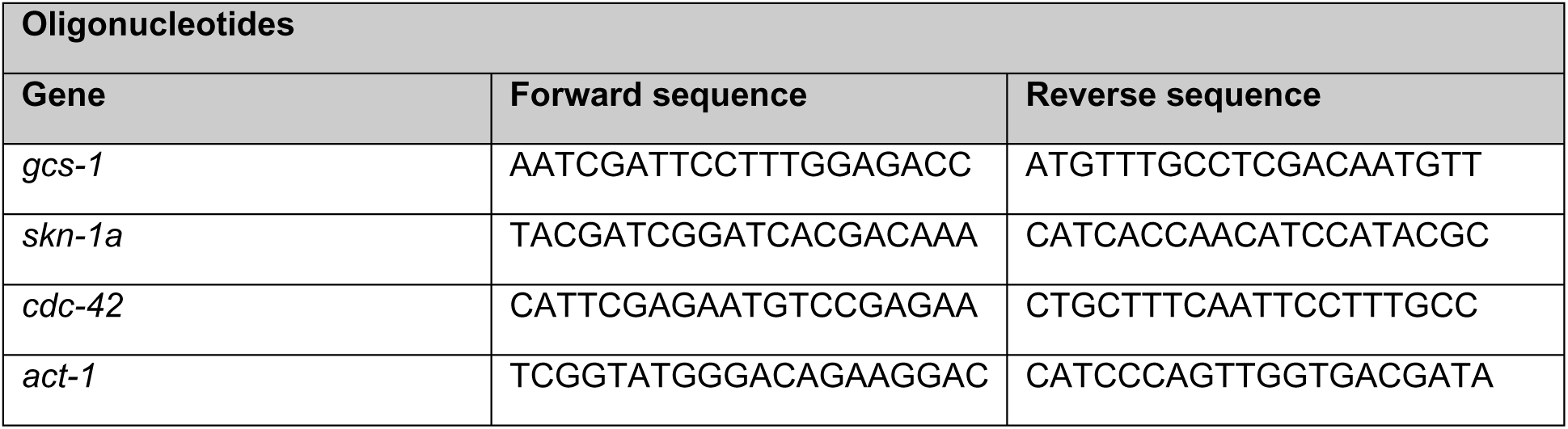

### RNAseq

Samples were prepared from ∼5000 synchronized, L1 arrested day-1 adult animals grown as described above for experiments involving GSC(-) animals. RNA was extracted using the same protocol described above for qRT-PCR samples. Purified RNA samples were DNase-treated and assigned an RNA Integrity Number (RIN) quality score using a Bioanalyzer 2100 (Agilent Technologies, Santa Clara, CA). Only matched samples with high RIN scores were sent for sequencing to Novogene Corporation Inc. (U.S. Subsidiary & UC Davis Sequencing Center). Briefly, for sequencing a total of 1 µg RNA per sample was used as input material for RNA sample preparations, then libraries were generated using NEBNext® Ultra TM RNA Library Prep Kit for Illumina® (NEB, USA) following the manufacturer’s recommendation, with index codes were added to attribute sequences to each sample. mRNA was purified from total RNA using poly-T oligo-attached magnetic beads, with fragmentation carried out using divalent cations under elevated temperature in NEBNext First Strand Synthesis Reaction Buffer (5X). First strand cDNA was synthesized using random hexamer primer and M-MuLV Reverse Transcriptase (RNase H-). Second strand cDNA synthesis was subsequently performed using DNA Polymerase I and RNase H. Remaining overhangs were converted into blunt ends via exonuclease/polymerase activities. After adenylation of 3’ ends of DNA fragments, NEBNext Adaptor with hairpin loop structure were ligated to prepare for hybridization. In order to select cDNA fragments of preferentially 150∼200 bp in length, the library fragments were purified with AMPure XP system (Beckman Coulter, Beverly, USA). Then 3 µl USER Enzyme (NEB, USA) was used with size-selected, adaptor-ligated cDNA at 37 °C for 15 min followed by 5 min at 95 °C before PCR. Then PCR was performed with Phusion High-Fidelity DNA polymerase, Universal PCR primers and Index (X) Primer. Finally, PCR products were purified (AMPure XP system) and library quality was assessed on the Agilent Bioanalyzer 2100 system. The clustering of the index-coded samples was performed on a cBot Cluster Generation System using PE Cluster Kit cBot- HS (Illumina) according to the manufacturer’s instructions.

After cluster generation, the library preparations were sequenced on an Illumina platform and paired-end reads were generated. Raw data (raw reads) of FASTQ format were firstly processed through fastp(*72*). In this step, clean data (clean reads) were obtained by removing reads containing adapter and poly-N sequences and reads with low quality from raw data. At the same time, Q20, Q30 and GC content of the clean data were calculated. All the downstream analyses were based on the clean data with high quality. Reference genome and gene model annotation files were downloaded from genome website browser (NCBI/UCSC/Ensembl) directly. Paired-end clean reads were mapped to the reference genome using HISAT2 software. HISAT2 uses a large set of small GFM indexes that collectively cover the whole genome. These small indexes (called local indexes), combined with several alignment strategies, enable rapid and accurate alignment of sequencing reads. Feature counts were used to count the read numbers mapped of each gene, including known and novel genes, then RPKM of each gene was calculated based on the length of the gene and reads count mapped to this gene. RPKM, Reads Per Kilobase of exon model per Million mapped reads, considers the effect of sequencing depth and gene length for the reads count at the same time, and is currently the most commonly used method for estimating gene expression levels(*73*). Differential expression analysis between two conditions/groups (three biological replicates per condition) was performed using DESeq2 R package(*74*). DESeq2 provides statistical routines for determining differential expression in digital gene expression data using a model based on the negative binomial distribution. The resulting P values were adjusted using the Benjamini and Hochberg’s approach for controlling the False Discovery Rate (FDR). Genes with an adjusted P value < 0.05 found by DESeq2 were assigned as differentially expressed. Gene categories were analyzed by the web-based tool WormCat(*75*).

### Proteomic analyses

We determined the overlap between our SKN-1 screen positives (including all 3 screens: genome-scale (suppressors and enhancers), phospholipid and LD) with the *C. elegans* LD proteome by analyzing two independent proteomic studies(*76, 77*). Both lists of proteins were curated for genes that had been named (or renamed) after these publications using WormBase.org. The lists were crossed using Venny2.1(*78*) and overlapping set of genes were analyzed on nemates.org (James Lund lab, University of Kentucky). Details of the lists and overlaps are deposited in **Supplementary Data 3**.

For overlap with the human liver LD proteome we converted all the peptides generated using the Huh7 cell line, a human hepatocellular carcinoma cell line(*79*) to *C. elegans* genes using WormBase.org and UniProt.org (see **Supplementary Data 3** for details). When a human peptide had more than one ortholog we used all potential *C. elegans* genes identified on WormBase.org or UniProt.org. The lists were crossed using Venny2.1(*78*) and overlapping set of genes were analyzed on nemates.org (James Lund lab, University of Kentucky).

## Acknowledgments

We thank Blackwell lab members for helpful discussions and critically reading the manuscript, and Peng Zhang for contributions during the project’s early stages.

## Funding

Supported by the:

American Federation for Aging Research (AFAR)/Glenn Foundation for Medical

Research Postdoctoral Fellowship (PD18019 to JIC-Q)

AFAR Reboot Fund (#REBOOT21004 to JIC-Q)

National Institutes of Health (AG54215 and GM122610 to TKB)

## Author contributions

Conceptualization: JICQ, TKB

Methodology: NJL

Investigation: JICQ, MJS, LPFC, NKP, ZW, FZ, NM, VT, MSB, NJL, LEMM, MT

Supervision: TKB

Writing—original draft: JICQ

Writing—review & editing: JICQ, TKB

## References

1. H. Shimano, R. Sato, SREBP-regulated lipid metabolism: Convergent physiology-divergent pathophysiology. Nat. Rev. Endocrinol. 13, 710–730 (2017).

2. J. Jacquemyn, A. Cascalho, R. E. Goodchild, The ins and outs of endoplasmic reticulum-controlled lipid biosynthesis. EMBO Rep. 18, 1905–1921 (2017).

3. H. Yki-Järvinen, P. K. Luukkonen, L. Hodson, J. B. Moore, Dietary carbohydrates and fats in nonalcoholic fatty liver disease. Nat. Rev. Gastroenterol. Hepatol. 18, 770–786 (2021).

4. D. D. Wang, F. B. Hu, Dietary Fat and Risk of Cardiovascular Disease: Recent Controversies and Advances. Annu. Rev. Nutr. 37, 423–446 (2017).

5. J. G. Wallis, J. L. Watts, J. Browse, Polyunsaturated fatty acid synthesis: What will they think of next? Trends Biochem. Sci. 27, 467–473 (2002).

6. T. C. Walther, J. Chung, R. V. R. V. Farese Jr., C. Daumer-Haas, S. Schuffenhauer, J. U. Walther, R. D. Schipper, T. Porstmann, J. R. Korenberg, T. C. Walther, J. Chung, R. V. R. V. Farese Jr., Lipid droplet biogenesis. Annu. Rev. Cell Dev. Biol. **33**, 491–510 (2017).

7. J. A. Olzmann, P. Carvalho, Dynamics and functions of lipid droplets. Nat. Rev. Mol. Cell Biol. 20, 137–155 (2019).

8. A. M. ALJohani, D. N. Syed, J. M. Ntambi, Insights into Stearoyl-CoA Desaturase-1 Regulation of Systemic Metabolism. Trends Endocrinol. Metab. 28, 831–842 (2017).

9. M. Hansen, T. Flatt, H. Aguilaniu, Reproduction, Fat Metabolism, and Life Span: What Is the Connection? Cell Metab. 17, 10–19 (2013).

10. V. Bustos, L. Partridge, Good Ol’ Fat: Links between Lipid Signaling and Longevity. Trends Biochem. Sci. 42, 812–823 (2017).

11. E. J. O’Rourke, A. A. Soukas, C. E. Carr, G. Ruvkun, C. elegans Major Fats Are Stored in Vesicles Distinct from Lysosome-Related Organelles. Cell Metab. 10, 430–435 (2009).

12. M. J. Steinbaugh, S. D. Narasimhan, S. Robida-Stubbs, L. E. Moronetti Mazzeo, J. M. Dreyfuss, J. M. Hourihan, P. Raghavan, T. N. Operaña, R. Esmaillie, T. K. Blackwell, Lipid-mediated regulation of SKN-1/Nrf in response to germ cell absence. Elife. 4, e07836 (2015).

13. Q. L. Wan, Z. L. Yang, X. G. Zhou, A. J. Ding, Y. Z. Pu, H. R. Luo, G. S. Wu, The effects of age and reproduction on the lipidome of Caenorhabditis elegans. Oxid. Med. Cell. Longev. 2019, 5768953 (2019).

14. J. Goudeau, S. Bellemin, E. Toselli-Mollereau, M. Shamalnasab, Y. Chen, H. Aguilaniu, Fatty acid desaturation links germ cell loss to longevity through NHR-80/HNF4 in C. elegans. PLoS Biol. 9, e1000599 (2011).

15. S. Han, E. A. Schroeder, C. G. Silva-García, K. Hebestreit, W. B. Mair, A. Brunet, Mono- unsaturated fatty acids link H3K4me3 modifiers to C. elegans lifespan. Nature. 544, 185–190 (2017).

16. S. Imanikia, M. Sheng, C. Castro, J. L. Griffin, R. C. Taylor, XBP-1 Remodels Lipid Metabolism to Extend Longevity. Cell Rep. 28, 581–589.e4 (2019).

17. T. K. Blackwell, M. J. Steinbaugh, J. M. Hourihan, C. Y. Ewald, M. Isik, SKN-1/Nrf, stress responses, and aging in Caenorhabditis elegans. Free Radic. Biol. Med. 88, 290–301 (2015).

18. N. J. Lehrbach, G. Ruvkun, Proteasome dysfunction triggers activation of SKN-1A/Nrf1 by the aspartic protease DDI-1. Elife. 5, e17721 (2016).

19. L. Baird, M. Yamamoto, The Molecular Mechanisms Regulating the KEAP1-NRF2 Pathway. Mol. Cell. Biol. 40, e00099–20 (2020).

20. A. Northrop, H. A. Byers, S. K. Radhakrishnan, Regulation of NRF1, a master transcription factor of proteasome genes: Implications for cancer and neurodegeneration. Mol. Biol. Cell. 31, 2158– 2163 (2020).

21. J. Steffen, M. Seeger, A. Koch, E. Krüger, Proteasomal degradation is transcriptionally controlled by TCF11 via an ERAD-dependent feedback loop. Mol. Cell. 40, 147–158 (2010).

22. N. Berner, K.-R. Reutter, D. H. Wolf, Protein Quality Control of the Endoplasmic Reticulum and Ubiquitin–Proteasome-Triggered Degradation of Aberrant Proteins: Yeast Pioneers the Path. Annu. Rev. Biochem. 87, 751–782 (2018).

23. J. A. Olzmann, R. R. Kopito, J. C. Christianson, The Mammalian Endoplasmic Reticulum- Associated Degradation System. Cold Spring Harb Perspect Biol. 5, a013185 (2015).

24. N. J. Lehrbach, G. Ruvkun, Endoplasmic reticulum-associated SKN- 1A/Nrf1 mediates a cytoplasmic unfolded protein response and promotes longevity. Elife. 8, e44425 (2019).

25. Z. Xu, L. Chen, L. Leung, T. S. B. Yen, C. Lee, J. Y. Chan, Liver-specific inactivation of the Nrf1 gene in adult mouse leads to nonalcoholic steatohepatitis and hepatic neoplasia. Proc. Natl. Acad. Sci. U. S. A. 102, 4120–5 (2005).

26. T. Tsujita, V. Peirce, L. Baird, Y. Matsuyama, M. Takaku, S. V. Walsh, J. L. Griffin, A. Uruno, M. Yamamoto, J. D. Hayes, Transcription Factor Nrf1 Negatively Regulates the Cystine/Glutamate Transporter and Lipid-Metabolizing Enzymes. Mol. Cell. Biol. 34, 3800–16 (2014).

27. X. Wu, T. A. Rapoport, Translocation of Proteins through a Distorted Lipid Bilayer. Trends Cell Biol. 31, 1–12 (2021).

28. J. H. An, T. K. Blackwell, SKN-1 links C. elegans mesendodermal specification to a conserved oxidative stress response. Genes Dev. 17, 1882–93 (2003).

29. D. Vilchez, I. Morantte, Z. Liu, P. M. Douglas, C. Merkwirth, A. P. C. Rodrigues, G. Manning, A. Dillin, RPN-6 determines C. elegans longevity under proteotoxic stress conditions. Nature. 489, 263–8 (2012).

30. J. L. Watts, M. Ristow, Lipid and carbohydrate metabolism in Caenorhabditis elegans. Genetics. 207, 413–446 (2017).

31. S. Pang, D. A. Lynn, J. Y. Lo, J. Paek, S. P. Curran, SKN-1 and Nrf2 couples proline catabolism with lipid metabolism during nutrient deprivation. Nat. Commun. 5, 5048 (2014).

32. J. D. Nhan, C. D. Turner, S. M. Anderson, C.-A. Yen, H. M. Dalton, H. K. Cheesman, D. L. Ruter, N. U. N. U. N. U. Naresh, C. M. Haynes, A. A. Soukas, R. Pukkila-Worley, S. P. Curran, Redirection of SKN-1 abates the negative metabolic outcomes of a perceived pathogen infection. Proc Natl Acad Sci U S A. 116, 22322–22330 (2019).

33. A. K. Walker, R. L. Jacobs, J. L. Watts, V. Rottiers, K. Jiang, D. M. Finnegan, T. Shioda, M. Hansen, F. Yang, L. J. Niebergall, D. E. Vance, M. Tzoneva, A. C. Hart, A. M. Näär, A conserved SREBP-1/phosphatidylcholine feedback circuit regulates lipogenesis in metazoans. Cell. 147, 840–852 (2011).

34. R. Panatala, H. Hennrich, J. C. M. Holthuis, Inner workings and biological impact of phospholipid flippases. J. Cell Sci. 128, 2021–2032 (2015).

35. A. Chorlay, L. Monticelli, J. Veríssimo Ferreira, K. Ben M’barek, D. Ajjaji, S. Wang, E. Johnson, R. Beck, M. Omrane, M. Beller, P. Carvalho, A. Rachid Thiam, Membrane Asymmetry Imposes Directionality on Lipid Droplet Emergence from the ER. Dev. Cell. 50, 25–42.e7 (2019).

36. S. Kim, C. Li, R. V. Farese, T. C. Walther, G. A. Voth, Key Factors Governing Initial Stages of Lipid Droplet Formation. J. Phys. Chem. B. 126, 453–462 (2022).

37. N. N. Lyssenko, Y. Miteva, S. Gilroy, W. Hanna-Rose, R. A. Schlegel, An unexpectedly high degree of specialization and a widespread involvement in sterol metabolism among the C. elegans putative aminophospholipid translocases. BMC Dev. Biol. 8, 96 (2008).

38. S. Cases, S. J. Stone, P. Zhou, E. Yen, B. Tow, K. D. Lardizabal, T. Voelker, R. V. Farese, Cloning of DGAT2, a Second Mammalian Diacylglycerol Acyltransferase, and Related Family Members. J. Biol. Chem. 276, 38870–38876 (2001).

39. N. Krahmer, Y. Guo, F. Wilfling, M. Hilger, S. Lingrell, K. Heger, H. W. Newman, M. Schmidt- Supprian, D. E. Vance, M. Mann, R. V. Farese, T. C. Walther, Phosphatidylcholine synthesis for lipid droplet expansion is mediated by localized activation of CTP:Phosphocholine cytidylyltransferase. Cell Metab. 14, 504–515 (2011).

40. M. Gao, X. Huang, B. L. Song, H. Yang, The biogenesis of lipid droplets: Lipids take center stage. Prog. Lipid Res. 75, 100989 (2019).

41. V. Choudhary, N. Ojha, A. Golden, W. A. Prinz, A conserved family of proteins facilitates nascent lipid droplet budding from the ER. J. Cell Biol. 211, 261–271 (2015).

42. K. Xie, P. Zhang, H. Na, Y. Liu, H. Zhang, P. Liu, MDT-28/PLIN-1 mediates lipid droplet- microtubule interaction via DLC-1 in Caenorhabditis elegans. Sci. Rep. 9, 14902 (2019).

43. E. L. Greer, T. J. Maures, A. G. Hauswirth, E. M. Green, D. S. Leeman, G. S. Maro, S. Han, M. R. Banko, O. Gozani, A. Brunet, Members of the H3K4 trimethylation complex regulate lifespan in a germline-dependent manner in C. elegans. Nature. 466, 383–7 (2010).

44. D. Bazopoulou, D. Knoefler, Y. Zheng, K. Ulrich, B. J. Oleson, L. Xie, M. Kim, A. Kaufmann, Y. Lee, Y. Dou, Developmental ROS individualizes organismal stress resistance and lifespan. Nature. 576, 301–305 (2019).

45. L. C. Tapsell, Foods and food components in the Mediterranean diet: Supporting overall effects. BMC Med. 12, 12–14 (2014).

46. Q. Gao, J. M. Goodman, The lipid droplet-a well-connected organelle. Front. Cell Dev. Biol. 3, 49 (2015).

47. J. Stevenson, E. Y. Huang, J. A. Olzmann, Endoplasmic Reticulum-Associated Degradation and Lipid Homeostasis. Annu. Rev. Nutr. 36, 511–542 (2016).

48. H. L. Ploegh, A lipid-based model for the creation of an escape hatch from the endoplasmic reticulum. Nature. 448, 435–438 (2007).

49. M. To, C. W. H. H. Peterson, M. A. Roberts, J. L. Counihan, T. T. Wu, M. S. Forster, D. K. Nomura, J. A. Olzmann, Lipid disequilibrium disrupts ER proteostasis by impairing ERAD substrate glycan trimming and dislocation. Mol. Biol. Cell. 28, 270–284 (2017).

50. M. T. Miedel, N. J. Graf, K. E. Stephen, O. S. Long, S. C. Pak, D. H. Perlmutter, G. A. Silverman, C. J. Luke, A pro-cathepsin L mutant is a luminal substrate for endoplasmic-reticulum-associated degradation in C. elegans. PLoS One. 7, e40145 (2012).

51. A. Mazzocchi, L. Leone, C. Agostoni, I. Pali-Schöll, The secrets of the mediterranean diet. Does [only] olive oil matter? Nutrients. 11, 2941 (2019).

52. A. Santinho, A. Chorlay, L. Foret, A. R. Thiam, Fat inclusions strongly alter membrane mechanics. Biophys. J. 120, 607–617 (2021).

53. J. Xu, S. Taubert, Beyond Proteostasis: Lipid metabolism as a new player in ER homeostasis. Metabolites. 11, 52 (2021).

54. M. A. Roberts, J. A. Olzmann, Protein Quality Control and Lipid Droplet Metabolism. Annu. Rev. Cell Dev. Biol. 36, 115–139 (2020).

55. V. Goder, E. Alanis-Dominguez, M. Bustamante-Sequeiros, Lipids and their (un)known effects on ER-associated protein degradation (ERAD). Biochim. Biophys. Acta - Mol. Cell Biol. Lipids. 1865, 158488 (2020).

56. G. Farrell, J. M. Schattenberg, I. Leclercq, M. M. Yeh, R. Goldin, N. Teoh, D. Schuppan, Mouse Models of Nonalcoholic Steatohepatitis: Toward Optimization of Their Relevance to Human Nonalcoholic Steatohepatitis. Hepatology. 69, 2241–2257 (2019).

57. S. B. Widenmaier, N. A. Snyder, T. B. Nguyen, A. Arduini, G. Y. Lee, A. P. Arruda, J. Saksi, A. Bartelt, G. S. Hotamisligil, NRF1 Is an ER Membrane Sensor that Is Central to Cholesterol Homeostasis. Cell. 171, 1094–1109.e15 (2017).

58. A. Silva-Palacios, M. Ostolga-Chavarría, C. Zazueta, M. Königsberg, Nrf2: Molecular and epigenetic regulation during aging. Ageing Res. Rev. 47, 31–40 (2018).

59. L. C. D. Pomatto, T. Dill, B. Carboneau, S. Levan, J. Kato, E. M. Mercken, K. J. Pearson, M. Bernier, R. de Cabo, Deletion of Nrf2 shortens lifespan in C57BL6/J male mice but does not alter the health and survival benefits of caloric restriction. Free Radic. Biol. Med. 152, 650–658 (2020).

60. S. Brenner, The genetics of Caenorhabditis elegans. Genetics. 77, 71–94 (1974).

61. P. Zhang, H.-Y. Qu, Z. Wu, H. Na, J. Hourihan, F. Zhang, F. Zhu, M. Isik, A. J. Walhout, Y.-X. Feng, T. K. Blackwell, bioRxiv, in press.

62. R. S. Kamath, J. Ahringer, Genome-wide RNAi screening in Caenorhabditis elegans. Methods. 30, 313–321 (2003).

63. J. F. Rual, J. Ceron, J. Koreth, T. Hao, A. S. Nicot, T. Hirozane-Kishikawa, J. Vandenhaute, S. H. Orkin, D. E. Hill, S. van den Heuvel, M. Vidal, Toward improving Caenorhabditis elegans phenome mapping with an ORFeome-based RNAi library. Genome Res. 14, 2162–2168 (2004).

64. C. Y. Ewald, J. N. Landis, J. P. Abate, C. T. Murphy, T. K. Blackwell, Dauer-independent insulin/IGF-1-signalling implicates collagen remodelling in longevity. Nature. 519, 97–101 (2015).

65. A. R. Turkish, S. L. Sturley, The genetics of neutral lipid biosynthesis: An evolutionary perspective. Am. J. Physiol. - Endocrinol. Metab. 297, 19–27 (2009).

66. N. Arantes-Oliviera, J. Apfeld, A. Dillin, C. Kenyon, N. Arantes-Oliveira, J. Apfeld, A. Dillin, C. Kenyon, Regulation of life-span by germ-line stem cells in Caenorhabditis elegans. Science. 295, 502–505 (2002).

67. D. H. Mitchell, J. W. Stiles, J. Santelli, D. Rao Sanadi, Synchronous growth and aging of Caenorhabditis elegans in the presence of fluorodeoxyuridine. Journals Gerontol. 34, 28–36 (1979).

68. J. I. Castillo-Quan, L. Li, K. J. Kinghorn, D. K. Ivanov, L. S. Tain, C. Slack, F. Kerr, T. Nespital, J. Thornton, J. Hardy, I. Bjedov, L. Partridge, Lithium Promotes Longevity through GSK3/NRF2- Dependent Hormesis. Cell Rep. 15, 638–650 (2016).

69. M. L. Deline, T. L. Vrablik, J. L. Watts, Dietary Supplementation of Polyunsaturated Fatty Acids in Caenorhabditis elegans. J. Vis. Exp. 81, e50879 (2013).

70. Y. Li, K. Na, H. J. Lee, E. Y. Lee, Y. K. Paik, Contribution of sams-1 and pmt-1 to lipid homoeostasis in adult Caenorhabditis elegans. J. Biochem. 149, 529–538 (2011).

71. X. Li, O. Matilainen, C. Jin, K. M. Glover-Cutter, C. I. Holmberg, T. K. Blackwell, Specific SKN- 1/Nrf stress responses to perturbations in translation elongation and proteasome activity. PLoS Genet. 7, e1002119 (2011).

72. P. J. A. Cock, C. J. Fields, N. Goto, M. L. Heuer, P. M. Rice, The Sanger FASTQ file format for sequences with quality scores, and the Solexa/Illumina FASTQ variants. Nucleic Acids Res. 38, 1767–1771 (2009).

73. C. Trapnell, B. A. Williams, G. Pertea, A. Mortazavi, G. Kwan, M. J. Van Baren, S. L. Salzberg, B. J. Wold, L. Pachter, Transcript assembly and quantification by RNA-Seq reveals unannotated transcripts and isoform switching during cell differentiation. Nat. Biotechnol. 28, 511–515 (2010).

74. S. Anders, W. Huber, Differential expression analysis for sequence count data. Genome Biol. 11, R106 (2010).

75. A. D. Holdorf, D. P. Higgins, A. C. Hart, P. R. Boag, G. J. Pazour, A. J. M. Walhout, A. K. Walker, WormCat: An online tool for annotation and visualization of caenorhabditis elegans genome-scale data. Genetics. 214, 279–294 (2020).

76. T. L. Vrablik, J. L. Watts, Polyunsaturated Fatty Acid Derived Signaling in Reproduction and Developmen: Insights From Caenorhabditis elegans and Drosophila melanogaster. Mol. Reprod. Dev. 80, 244–259 (2013).

77. P. Zhang, H. Na, Z. Liu, S. S. O. S. Zhang, P. Xue, Y. Chen, J. Pu, G. Peng, X. Huang, F. Yang, Z. Xie, T. Xu, P. Xu, G. Ou, S. S. O. S. Zhang, P. Liu, Proteomic Study and Marker Protein Identification of Caenorhabditis elegans Lipid Droplets. Mol. Cell. Proteomics. 11, 317–328 (2012).

78. 78. J. C. Oliveiros, Venny. An interactive tool for comparing lists with Venn’s diagrams. (2015), p. https://bioinfogp.cnb.csic.es/tools/venny/index.ht, (available at https://bioinfogp.cnb.csic.es/tools/venny/index.html).

79. K. Bersuker, C. W. H. Peterson, M. To, S. J. Sahl, V. Savikhin, E. A. Grossman, D. K. Nomura, J. Olzmann, A Proximity Labeling Strategy Provides Insights into the Composition and Dynamics of Lipid Droplet Proteomes. Dev. Cell. 44, 97–112.e7 (2018).

## Supplementary References

80. N. J. Lehrbach, P. C. Breen, G. Ruvkun, Protein Sequence Editing of SKN-1A/Nrf1 by Peptide:N- Glycanase Controls Proteasome Gene Expression. Cell. 177, 737–750 (2019).

81. T. L. Vrablik, V. A. Petyuk, E. M. Larson, R. D. Smith, J. L. Watts, Lipidomic and proteomic analysis of Caenorhabditis elegans lipid droplets and identification of ACS-4 as a lipid droplet- associated protein. Biochim. Biophys. Acta - Mol. Cell Biol. Lipids. 1851, 1337–1345 (2015).

82. N. Bodnar, T. Rapoport, Toward an understanding of the Cdc48/p97 ATPase. F1000Research. 6, 1318 (2017).

83. J. Mouysset, C. Kähler, T. Hoppe, A conserved role of Caenorhabditis elegans CDC-48 in ER- associated protein degradation. J. Struct. Biol. 156, 41–49 (2006).

84. J. Pispa, S. Palmén, C. I. Holmberg, J. Jäntti, C. elegans dss-1 is functionally conserved and required for oogenesis and larval growth. BMC Dev. Biol. 8, 1–13 (2008).

85. C. Schuberth, A. Buchberger, UBX domain proteins: Major regulators of the AAA ATPase Cdc48/p97. Cell. Mol. Life Sci. 65, 2360–2371 (2008).

86. R. Sonneville, S. P. Moreno, A. Knebel, C. Johnson, C. J. Hastie, A. Gartner, A. Gambus, K. Labib, CUL-2LRR-1 and UBXN-3 drive replisome disassembly during DNA replication termination and mitosis. Nat. Cell Biol. 19, 468–479 (2017).

87. J. A. Olzmann, C. M. Richter, R. R. Kopito, Spatial regulation of UBXD8 and p97/VCP controls ATGL-mediated lipid droplet turnover. Proc. Natl. Acad. Sci. 110, 1345–1350 (2013).

88. J. N. Lee, H. Kim, H. Yao, Y. Chen, K. Weng, J. Ye, Identification of Ubxd8 protein as a sensor for unsaturated fatty acids and regulator of triglyceride synthesis. Proc. Natl. Acad. Sci. U. S. A. 107, 21424–9 (2010).

89. M. A. Surma, C. Klose, D. Peng, M. Shales, C. Mrejen, A. Stefanko, H. Braberg, D. E. Gordon, D. Vorkel, C. S. Ejsing, R. Farese, K. Simons, N. J. Krogan, R. Ernst, A lipid E-MAP identifies Ubx2 as a critical regulator of lipid saturation and lipid bilayer stress. Mol. Cell. 51, 519–530 (2013).

90. Y. Sasagawa, K. Yamanaka, T. Ogura, ER E3 ubiquitin ligase HRD-1 and its specific partner chaperone BiP play important roles in ERAD and developmental growth in Caenorhabditis elegans. Genes to Cells. 12, 1063–1073 (2007).

91. J. Schulz, D. Avci, M. A. Queisser, A. Gutschmidt, L. S. Dreher, E. J. Fenech, N. Volkmar, Y. Hayashi, T. Hoppe, J. C. Christianson, Conserved cytoplasmic domains promote Hrd1 ubiquitin ligase complex formation for ER-associated degradation (ERAD). J. Cell Sci. 130, 3322–3335 (2017).

92. M. Zhen, J. E. Schein, D. L. Baillie, E. Peter, M. Candido, An essential ubiquitin-conjugating enzyme with tissue and developmental specificity in the nematode Caenorhabditis elegans. EMBO J. 15, 3229–3237 (1996).

93. M. J. Kinet, J. A. Malin, M. C. Abraham, E. S. Blum, M. R. Silverman, Y. Lu, S. Shaham, HSF-1 activates the ubiquitin proteasome system to promote non-apoptotic developmental cell death in C. elegans. Elife. 5, 1–24 (2016).

94. P. Gupta, L. Leahul, X. Wang, C. Wang, B. Bakos, K. Jasper, D. Hansen, Proteasome regulation of the chromodomain protein MRG-1 controls the balance between proliferative fate and differentiation in the C. elegans germ line. Development. 142, 291–302 (2015).

95. E. Crowe, E. P. M. Candido, Characterization of C. elegans RING finger protein 1, a binding partner of ubiquitin-conjugating enzyme 1. Dev. Biol. 265, 446–459 (2004).

96. A. Davy, P. Bello, N. Thierry-Mieg, P. Vaglio, J. Hitti, L. Doucette-Stamm, D. Thierry-Mieg, J. Reboul, S. Boulton, A. J. M. Walhout, O. Coux, M. Vidal, A protein-protein interaction map of the Caenorhabditis elegans 26S proteasome. EMBO Rep. 2, 821–828 (2001).

97. F. Urano, M. Calfon, T. Yoneda, C. Yun, M. Kiraly, S. G. Clark, D. Ron, A survival pathway for Caenorhabditis elegans with a blocked unfolded protein response. J. Cell Biol. 158, 639–646 (2002).

98. S. Yamauchi, Y. Sasagawa, T. Ogura, K. Yamanaka, Differential expression pattern of UBX family genes in Caenorhabditis elegans. Biochem. Biophys. Res. Commun. 358, 545–552 (2007).

99. L. J. Smulan, W. Ding, E. Freinkman, S. Gujja, Y. J. K. Edwards, A. K. Walker, Cholesterol- Independent SREBP-1 Maturation Is Linked to ARF1 Inactivation. Cell Rep. 16, 9–18 (2016).

100. B. Chen, Y. Jiang, S. Zeng, J. Yan, X. Li, Y. Zhang, W. Zou, X. Wang, Endocytic sorting and recycling require membrane phosphatidylserine asymmetry maintained by TAT-1/CHAT-1. PLoS Genet. 6, e1001235 (2010).

101. B. He, J. Zhang, Y. Wang, Y. Li, X. Zou, B. Liang, Identification of cytochrome b5 CYTB-5.1 and CYTB-5.2 in C. elegans; evidence for differential regulation of SCD. Biochim. Biophys. Acta - Mol. Cell Biol. Lipids. 1863, 235–246 (2018).

102. Y. Zhang, H. Wang, J. Zhang, Y. Hu, L. Zhang, X. Wu, X. Su, T. Li, X. Zou, B. Liang, The cytochrome b5 reductase HPO-19 is required for biosynthesis of polyunsaturated fatty acids in Caenorhabditis elegans. Biochim. Biophys. Acta - Mol. Cell Biol. Lipids. 1861, 310–319 (2016).

103. S. Hashmi, Y. Wang, R. S. Parhar, K. S. Collison, W. Conca, F. Al-Mohanna, R. Gaugler, A C. elegans model to study human metabolic regulation. Nutr. Metab. 10, 1–11 (2013).

104. A. C. Carrano, A. Dillin, T. Hunter, A krüppel-like factor downstream of the E3 ligase WWP-1 mediates dietary-restriction-induced longevity in caenorhabditis elegans. Nat. Commun. 5 (2014), doi:10.1038/ncomms4772.

105. S. Taubert, M. R. Van Gilst, M. Hansen, K. R. Yamamoto, A mediator subunit, MDT-15, integrates regulation of fatty acid metabolism by NHR-49-dependent and -independent pathways in C. elegans. Genes Dev. 20, 1137–1149 (2006).

106. M. R. Van Gilst, H. Hadjivassiliou, A. Jolly, K. R. Yamamoto, Nuclear hormone receptor NHR-49 controls fat consumption and fatty acid composition in C. elegans. PLoS Biol. 3, 0301–0312 (2005).

107. D. Palgunow, M. Klapper, F. Döring, Dietary Restriction during Development Enlarges Intestinal and Hypodermal Lipid Droplets in Caenorhabditis elegans. PLoS One. 7 (2012), doi:10.1371/journal.pone.0046198.

108. P. P. Pathare, A. Lin, K. E. Bornfeldt, S. Taubert, M. R. van Gilst, Coordinate regulation of lipid metabolism by novel nuclear receptor partnerships. PLoS Genet. 8 (2012), doi:10.1371/journal.pgen.1002645.

109. H. Thieringer, B. Moellers, G. Dodt, W. H. Kunau, M. Driscoll, Modeling human peroxisome biogenesis disorders in the nematode Caenorhabditis elegans. J. Cell Sci. 116, 1797–1804 (2003).

110. J. Kong, Y. Ji, Y. G. Jeon, J. S. Han, K. H. Han, J. H. Lee, G. Lee, H. Jang, S. S. Choe, M. Baes, J. B. Kim, Spatiotemporal contact between peroxisomes and lipid droplets regulates fasting- induced lipolysis via PEX5. Nat. Commun. 11, 1–16 (2020).

111. M. Hanna, L. Wang, A. Audhya, Worming Our Way In and Out of the Caenorhabditis elegans Germline and Developing Embryo. Traffic. 14, 471–478 (2013).

112. H. W. Shin, H. Takatsu, Substrates of P4-ATPases: beyond aminophospholipids (phosphatidylserine and phosphatidylethanolamine). FASEB J. 33, 3087–3096 (2019).

113. M. D. Kilwein, M. A. Welte, Lipid Droplet Motility and Organelle Contacts. *Contact*. **2**, 251525641989568 (2019).

114. A. L. Schuh, M. Hanna, K. Quinney, L. Wang, A. Sarkeshik, J. R. Yates, A. Audhya, The VPS-20 subunit of the endosomal sorting complex ESCRT-III exhibits an open conformation in the absence of upstream activation. Biochem. J. 466, 625–637 (2015).

115. L. Luo, M. Hannemann, S. Koenig, J. Hegermann, M. Ailion, M. K. Cho, N. Sasidharan, M. Zweckstetter, S. A. Rensing, S. Eimer, The Caenorhabditis elegans GARP complex contains the conserved Vps51 subunit and is required to maintain lysosomal morphology. Mol. Biol. Cell. 22, 2564–2578 (2011).

116. O. Billing, B. Natarajan, A. Mohammed, P. Naredi, G. Kao, A directed RNAi screen based on larval growth arrest reveals new modifiers of C. elegans insulin signaling. PLoS One. 7, e34507 (2012).

117. F. Wilfling, A. R. Thiam, M. J. Olarte, J. Wang, R. Beck, T. J. Gould, E. S. Allgeyer, F. Pincet, J. Bewersdorf, R. V. Farese, T. C. Walther, Arf1/COPI machinery acts directly on lipid droplets and enables their connection to the ER for protein targeting. Elife. 2014, 1–20 (2014).

118. M. Zerial, H. McBride, Rab proteins as membrane organizers. Nat. Rev. Mol. Cell Biol. 2, 107–117 (2001).

119. P. Liu, R. Bartz, J. K. Zehmer, Y. shu Ying, M. Zhu, G. Serrero, R. G. W. Anderson, Rab-regulated interaction of early endosomes with lipid droplets. Biochim. Biophys. Acta - Mol. Cell Res. 1773, 784–793 (2007).

120. R. Bartz, J. K. Zehmer, M. Zhu, Y. Chen, G. Serrero, Y. Zhao, P. Liu, Dynamic activity of lipid droplets: Protein phosphorylation and GTP-mediated protein translocation. J. Proteome Res. 6, 3256–3265 (2007).

121. K. M. Brendza, W. Haakenson, R. E. Cahoon, L. M. Hicks, L. H. Palavalli, B. J. Chiapelli, M. McLaird, J. P. McCarter, D. J. Williams, M. C. Hresko, J. M. Jez, Phosphoethanolamine *N* - methyltransferase (PMT-1) catalyses the first reaction of a new pathway for phosphocholine biosynthesis in *Caenorhabditis elegans*. Biochem. J. 404, 439–448 (2007).

122. L. H. Palavalli, K. M. Brendza, W. Haakenson, R. E. Cahoon, M. McLaird, L. M. Hicks, J. P. McCarter, D. J. Williams, M. C. Hresko, J. M. Jez, Defining the role of phosphomethylethanolamine N-methyltransferase from Caenorhabditis elegans in phosphocholine biosynthesis by biochemical and kinetic analysis. Biochemistry. 45, 6056–6065 (2006).

123. J. H. Koh, L. Wang, C. Beaudoin-Chabot, G. Thibault, Lipid bilayer stress-activated IRE-1 modulates autophagy during endoplasmic reticulum stress. J. Cell Sci. 131, jcs217992 (2018).

124. J. Deng, X. Bai, H. Tang, S. Pang, DNA damage promotes ER stress resistance through elevation of unsaturated phosphatidylcholine in Caenorhabditis elegans. J. Biol. Chem. 296, 100095 (2021).

125. S. Li, S. Xu, Y. Ma, S. Wu, Y. Feng, Q. Cui, L. Chen, S. Zhou, Y. Kong, X. Zhang, J. Yu, M. Wu, S. O. Zhang, A Genetic Screen for Mutants with Supersized Lipid Droplets in Caenorhabditis elegans. G3 (Bethesda, Md.) Genes|Genomes|Genetics. 6, 2407–2419 (2016).

126. S. O. Zhang, A. C. Box, N. Xu, J. Le Men, J. Yu, F. Guo, R. Trimble, H. Y. Mak, Genetic and dietary regulation of lipid droplet expansion in Caenorhabditis elegans. Proc. Natl. Acad. Sci. U. S. A. 107, 4640–4645 (2010).

127. A. R. Kimmel, C. Sztalryd, The Perilipins: Major Cytosolic Lipid Droplet–Associated Proteins and Their Roles in Cellular Lipid Storage, Mobilization, and Systemic Homeostasis. Annu. Rev. Nutr. 36, 471–509 (2016).

128. H. Na, P. Zhang, Y. Chen, X. Zhu, Y. Liu, Y. Liu, K. Xie, N. Xu, F. Yang, Y. Yu, S. Cichello, H. Y. Mak, M. C. Wang, H. Zhang, P. Liu, Identification of lipid droplet structure-like/resident proteins in Caenorhabditis elegans. Biochim. Biophys. Acta - Mol. Cell Res. 1853, 2481–2491 (2015).

